# Distinct silencer states determine epigenetic states of heterochromatin

**DOI:** 10.1101/2022.02.01.478725

**Authors:** Daniel S. Saxton, Jasper Rine

## Abstract

A remarkable property of heterochromatin is that a given heterochromatic locus can exhibit different transcriptional states in genetically identical cells. Studies demonstrate that epigenetic inheritance of the silenced state requires silencers and therefore cannot be driven by the inheritance of modified histones alone. To address these observations, we surveyed the chromatin architectures of strong and weak silencers in *Saccharomyces*. We found that strong silencers recruited Sir proteins and silenced the locus in all cells. Strikingly, weakening these silencers reduced Sir protein recruitment and stably silenced the locus in some cells; however, this silenced state could probabilistically convert to an expressed state that lacked Sir protein recruitment. Additionally, changes in the constellation of silencer-bound proteins or the concentration of a structural Sir protein modulated the probability that a locus existed in the silenced or expressed state. These findings argued that distinct states of silencers generate epigenetic states and regulate their dynamics.

## Introduction

The ability of genetically identical cells to exhibit different transcriptional states is crucial both for multicellular organisms and for unicellular organisms that can assume multiple identities. In many cases, these transcriptional states can be transmitted to daughter cells and thereby constitute a type of memory that is epigenetic, or “above the genome”. The inheritance of some transcriptional states is clearly linked to mechanisms like DNA methylation, RNA interference, and transcription factor feedback loops. The transmission of modified histones through DNA replication provides an additional mechanism that could drive epigenetic memory, especially in the case of heterochromatic silencing. Interestingly, although this histone-based memory model is widely embraced, the degree to which it drives *bona fide* transcriptional memory remains unclear and controversial.

To establish a silenced heterochromatin state, silencing proteins bind to silencers, spread across adjacent nucleosomes, and modify histones. During DNA replication, daughter chromatids locally inherit modified histone H3-H4 tetramers from the parental chromatid and receive newly synthesized histones from the nucleoplasm (Jackson, 1988; Prior et al., 1980; Schlissel and Rine, 2019). A remarkable property of many silencing proteins is that they can both modify histones and bind to the modifications they generate (Hansen et al., 2008; Hecht et al., 1995; Imai et al., 2000; Zhang et al., 2008). As such, the histone-based memory model posits that silencing proteins bind to modified histones on daughter chromatids and modify adjacent, naïve histones to maintain the silenced state. Therefore, epigenetic memory could be driven by the combination of modified histone inheritance and a positive feedback loop between histone modifications and silencing protein recruitment.

The histone-based memory model makes multiple predictions that have been previously tested in *S. cerevisiae, S. pombe*, and *Drosophila*. First, mutations that reduce the inheritance of H3-H4 tetramers to daughter chromatids should lead to defects in silencing inheritance. However, mutations in two replication-coupled histone chaperones, Dpb3 and Mcm2, lead to severe reductions in histone inheritance and only minor defects in silencing inheritance (Saxton and Rine, 2019; Schlissel and Rine, 2019). Second, if histones were sufficient to carry the memory of the silenced state, then silencers would be dispensable for memory once the silenced state is established. However, the induced removal of silencer activity also removes the ability of a silenced state to be subsequently inherited, demonstrating that inheritance requires the constant presence of silencers and is not driven by modified histones alone (Audergon et al., 2015; Coleman and Struhl, 2017; Holmes and Broach, 1996; Laprell et al., 2017; Ragunathan et al., 2015). Thus, multiple studies argue that the histone-based memory model is either incorrect or incomplete. (A notable exception, revealed by the *epe1Δ* mutation in *S. pombe*, is discussed below.)

Studies of *bona fide* silencing inheritance require the existence of genetically identical cells that are heritably silenced or expressed at a locus of interest. Though many studies utilize these epigenetic states to study epigenetic inheritance, less attention has been given to the question of why some cells exhibit silencing and others do not. Clearly, one or more pivotal determinants of the silenced state can be active in one cell but inactive in another, genetically identical cell. Furthermore, the ability of a cell with a silenced locus to transmit the silenced state of that locus to daughter cells may be driven by the heritable activity of these determinants. Previous studies have partially characterized these determinants with a *cis*-trans test, in which two alleles of a gene co-exist within one cell, and both alleles can be either expressed or silenced. These studies found that the silenced state can exist at one allele while the other remains expressed, demonstrating that these determinants act in *cis* at each locus (Berry et al., 2015; Grewal and Klar, 1996; Xu et al., 2006). However, the identity of these *cis*-acting determinants of the silenced state remains unclear.

Much is known about heterochromatic silencing in the budding yeast *Saccharomyces cerevisiae.* Budding yeast have two mating-type loci, *HML* and *HMR*, that are silenced by the action of silencers and four Sir proteins, Sir1-4 (Rine and Herskowitz, 1987). To establish silencing, Sir proteins are recruited to the *E* and *I* silencers that each flank *HML* and *HMR*. This process is orchestrated by combinations of the DNA binding proteins ORC, Rap1, and Abf1, which bind in different combinations at each silencer and interact with Sir proteins. Sir1 binds only at silencers via an interaction with ORC, whereas Sir2-4 form the Sir complex which binds at silencers via interactions with Sir1, Rap1, and Abf1. Additionally, the ability of the Sir complex to deacetylate histones and bind efficiently to deacetylated histones allows it to spread from silencers across adjacent nucleosomes (Hoppe et al., 2002; Rusché et al., 2002; Thurtle and Rine, 2014). This form of silencing is not affected by DNA methylation or RNAi, as these mechanisms do not exist in budding yeast.

Both *HML* and *HMR* are constitutively silenced, and removal of Sir2, Sir3, or Sir4 leads to full expression of both loci (Rine and Herskowitz, 1987). In contrast, removal of Sir1 causes genetically identical cells to be either completely silenced or completely expressed at either locus (Dodson and Rine, 2015; Pillus and Rine, 1989; Xu et al., 2006). These different states are robustly transmitted to daughter cells, though rare events can occur in which a silenced state switches to an expressed state, or vice versa. Therefore, *sir1Δ* generates *bona fide* epigenetic states at *HML* and *HMR.* What are the determinants of the silenced state in *sir1Δ?* We reasoned that since these determinants act in *cis*, they should exist at a locus, such as *HML* or *HMR*, when that locus is silenced but not when it is expressed. To test this, we used Chromatin Immunoprecipitation (ChIP) to assess the chromatin architecture of different epigenetic states of *HML* and *HMR* in *sir1Δ* cells. Upon finding candidate determinants of the silenced state, we then modulated the activity of the determinants and measured the effects on epigenetic states.

## Results

### Silencer activity was bistable in *sir1Δ*

To identify *cis*-acting determinants of the silenced state, we used Fluorescence Activated Cell Sorting (FACS) to sort *sir1Δ* cells that exhibited different epigenetic states, and ChIP to survey the chromatin architecture in these different populations. To facilitate these experiments, we utilized versions of *HML* and *HMR* that encode fluorescent reporters expressed from the native *HML α* promoter: *HMLα::RFP* and *HMRα::GFP* (Figure 1A) (Saxton and Rine, 2019). Additionally, we replaced endogenous *SIR3* or *SIR4* with epitope-tagged versions of these genes (*SIR3-myc* or *SIR4-myc*). Consistent with previous studies, *HMLα::RFP* and *HMRα::GFP* were fully silenced in *SIR^+^* cells, fully expressed in *sir4Δ* cells, and either fully silenced or fully expressed in *sir1Δ* cells (Figure 1A) (Pillus and Rine, 1989; Saxton and Rine, 2019; Xu et al., 2006). We utilized FACS to sort *sir1Δ* cells into separated populations of cells that were silenced or expressed at *HMLα::RFP;* similarly, we sorted *sir1Δ* cells that were silenced or expressed at *HMRα::GFP* (Figure 1B). Because these sorted populations were not large enough for ChIP assays, we allowed these populations to double approximately eight times immediately after sorting. During this time, a small fraction of cells in each population switched to the opposite state; however, the final percentage of cells with the sorted state was always >70% and usually ~90% (Figure 1C-J). Finally, control strains *SIR^+^* and *sir4Δ* were not sorted but otherwise subjected to the same growth regimen as sorted *sir1Δ* cells.

**Figure 1:**
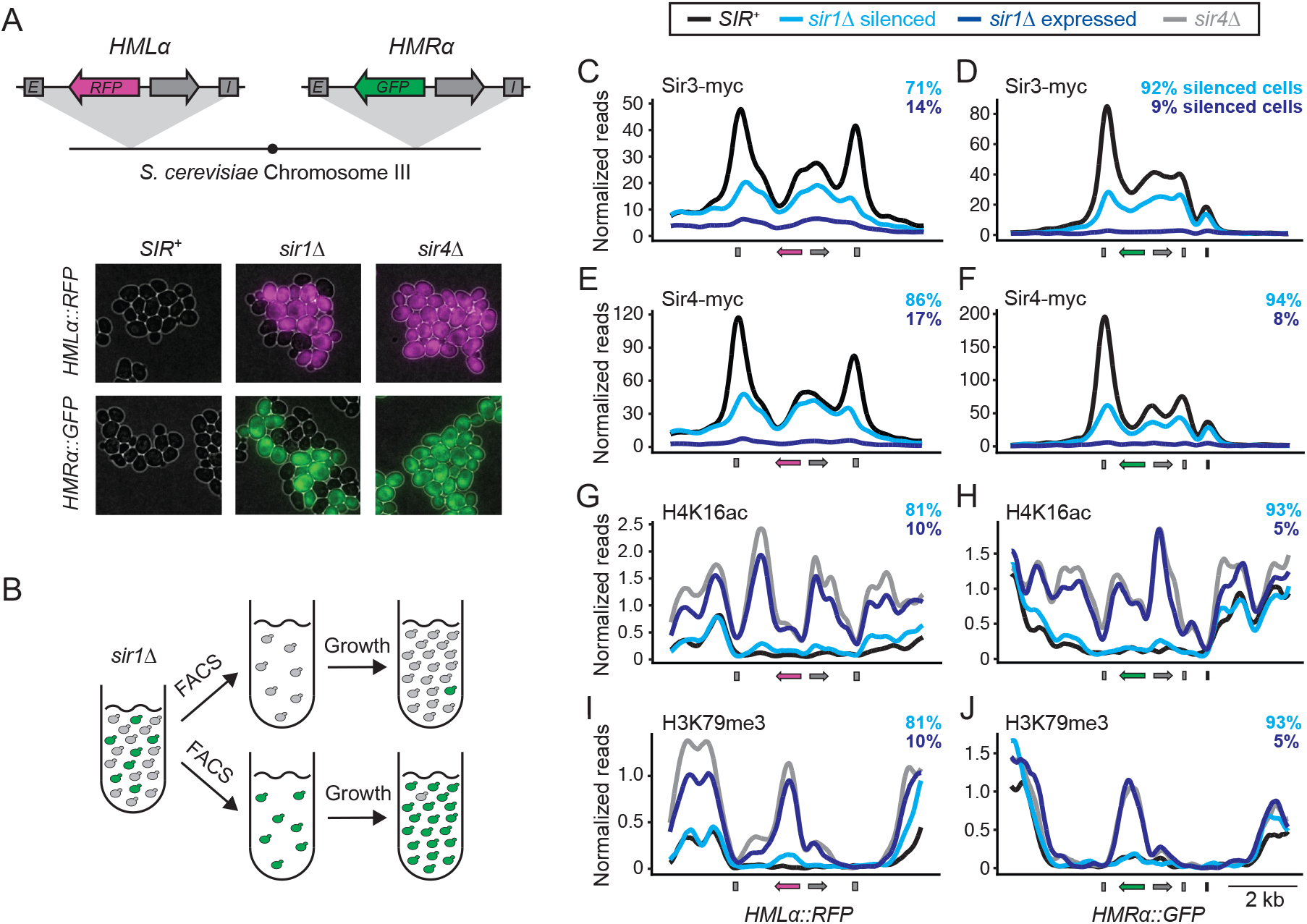
Silencers were bistable in *sir1Δ* cells. A. Schematic of *HMLα::RFP* and *HMRα::GFP*. Each locus contains *E* and *I* silencers (grey boxes), the *α1* gene (grey arrows), a fluorescent reporter gene (magenta or green arrows), and the bidirectional *α* promoter (located between both genes). Silencing phenotypes of either *HMLα::RFP* or *HMRα::GFP* in *SIR^+^, sir4Δ*, or *sir1Δ* cells are shown (JRY11494, JRY11496, and JRY13129-13132). B. Schematic of cell sorting and growth regimen for strains containing *HMRα::GFP.* C-J. ChIP of different Sir proteins and histone modifications in sorted and control populations. All strains contained either *SIR3-myc* or *SIR4-myc*, and either *HMLα::RFP* or *HMRα::GFP* (JRY11494, JRY11496, and JRY13125-13132). The light blue and dark blue percentages in each panel reflect the percentages of silenced cells for different sorted *sir1Δ* populations, as calculated by flow cytometry. Genomic features are shown at the bottom of each panel: grey boxes are silencers, arrows are genes, and the black box next to *HMRα::GFP* is a tRNA gene that exhibits a low degree of Sir protein binding. Reads were normalized to the genome-wide median. For C-F, an anti-myc antibody was used to assess Sir3-myc and Sir4-myc binding at silenced loci. E-J contain data from strains with *SIR4-myc*, but anti-H4K16ac and anti-H3K79me3 antibodies were used to assess the status of these histone modifications. The percentage of silenced cells were identical between G and I because chromatin for each was prepared from common cell populations and split for ChIP with different antibodies. This procedure also accounted for the similar numbers of silenced cells in H and J. Biological replicates, quantification, and additional ChIP data from other conditions are shown in Figures S1-S2.

Consistent with past ChIP studies, Sir3-myc and Sir4-myc localized robustly to both *HMLα::RFP* and *HMRα::GFP* in *SIR^+^* cells (Figure 1C-F, Figures S1-S2) (Hoppe et al., 2002; Rusché et al., 2002; Thurtle and Rine, 2014). Prominent binding peaks for both proteins were observed at the *E* and *I* silencers. An additional peak was observed at the *α* promoter, which has a Rap1 binding site that is necessary for transcription but also contributes to silencing (Cheng and Gartenberg, 2000; Siliciano and Tatchell, 1984). Strikingly, the *sir1Δ* silenced populations exhibited a large reduction in binding of Sir3-myc and Sir4-myc at silencers, and a modest reduction in binding at the *HMRα* promoter. In contrast, the *sir1Δ* expressed population exhibited no measurable binding of Sir3-myc or Sir4-myc at silencers or the *α* promoter at either locus. Importantly, Sir3-myc and Sir4-myc expression levels were similar between *SIR^+^* and *sir1Δ*, as evaluated by western blot (Figure S2P). Therefore, in *sir1Δ*, the silenced states of *HMLα::RFP* and *HMRα::GFP* exhibited reduced Sir complex binding and the expressed states of these loci lacked detectable Sir complex binding.

We next asked whether histone modifications characteristic of silenced chromatin were present in these different populations. Sir2 is the member of the Sir complex that deacetylates histones, and a notable example is the deacetylation of H4K16 (Imai et al., 2000; Landry et al., 2000). Additionally, silencing requires the removal of H3K79me3, a process that is driven by histone turnover during DNA replication (Leeuwen et al., 2002; Vos et al., 2011). As seen in previous studies, *SIR^+^* cells exhibited deacetylated H4K16 and unmethylated H3K79 at both *HMLα::RFP* and *HMRα::GFP* (Figure 1G-J, Figures S1-S2) (Hoppe et al., 2002; Ng et al., 2003; Rusché et al., 2002). In contrast, *sir4Δ* cells had H4K16ac and H3K79me3 at these loci.

Surprisingly, even though the *sir1Δ* silenced population exhibited reduced Sir complex binding, H4K16 was deacetylated and H3K79 was unmethylated to similar degrees as in *SIR^+^* cells. Consistent with the absence of Sir complex binding in the *sir1Δ* expressed population, this population showed similar levels of H4K16ac and H3K79me3 as *sir4Δ*.

Together, these findings argued that in *sir1Δ* cells, reduced Sir complex binding at silencers generated a destabilized version of heterochromatin that could spontaneously convert to a euchromatic state. This idea is consistent with the role of Sir1, which physically interacts with both silencer-bound ORC and Sir4, yet has no other known silencing function (Gardner et al., 1999; Triolo and Sternglanz, 1996). Additionally, the sorted *sir1Δ* silenced and expressed populations showed entirely different molecular architectures: Sir proteins partially nucleated at silencers, partially spread, and fully modified histones in cells with the silenced state at *HMLα::RFP* or *HMRα::GFP*, yet none of these processes occurred when these loci exhibited the expressed state. Given that nucleation (the recruitment of Sir proteins to silencers) is required for Sir protein spreading and modification of histones, our data argued that one pivotal determinant of these epigenetic states is the bistable binding of Sir proteins at silencers.

In theory, epigenetic states could emerge from cooperative binding of the Sir complex and feedback between the Sir complex and histone modifications (Sneppen and Dodd, 2015). In this model, the abilities of the Sir complex to bind to silencers, bind to nucleosomes, modify histones, and stabilize additional Sir complex binding would be probabilistic events that reinforce each other. The successful formation and maintenance of this feedback loop in some *sir1Δ* cells, but not others, could produce different epigenetic states. We designed a series of experiments to test this cooperativity model. Given that the silenced domain of *HMLα::RFP* extends to a nearby telomere and the silenced domain of *HMRα::GFP* does not (Ellahi et al., 2015; Thurtle and Rine, 2014), we performed most additional experiments at *HMRα::GFP*.

### Sir4 dosage determined the probability of silencing

If Sir complex binding at silencers and nucleosomes were cooperative, probabilistic events, we reasoned that changing the dosage of a Sir complex member may alter these probabilities and thus alter the frequency of silenced and expressed cells in *sir1Δ*. A previous study found that overexpression of either Sir3 or Sir4 partially rescues silencing defects observed in *sir1Δ*, as measured by population-level silencing assays (Dhillon and Kamakaka, 2000). We tested this question at the single-cell level by asking whether incremental increases or reductions in Sir4 dosage altered the frequency of epigenetic states in *sir1Δ* cells. To change Sir4 dosage, we used a heterologous fusion protein containing the DNA-binding protein LexA, the estradiol-binding domain of the human estrogen receptor (ER), and an *E. coli*-derived Activating Domain (AD) (Figure 2A). In addition, we replaced the endogenous *SIR4* promoter with a heterologous promoter containing a single *LexO* site. The addition of estradiol allows the LexA-ER-AD fusion protein to bind to the *LexO* site and activate transcription of a target gene (Ottoz et al., 2014). Similarly, we found that addition of estradiol to a strain containing *LexA-ER-AD* and *LexO-SIR4* led to expression of Sir4 and silencing of *HMRα::GFP*, and that the addition of estradiol did not affect silencing in cells lacking these inducible promoter components (Figure S3A-B).

**Figure 2:**
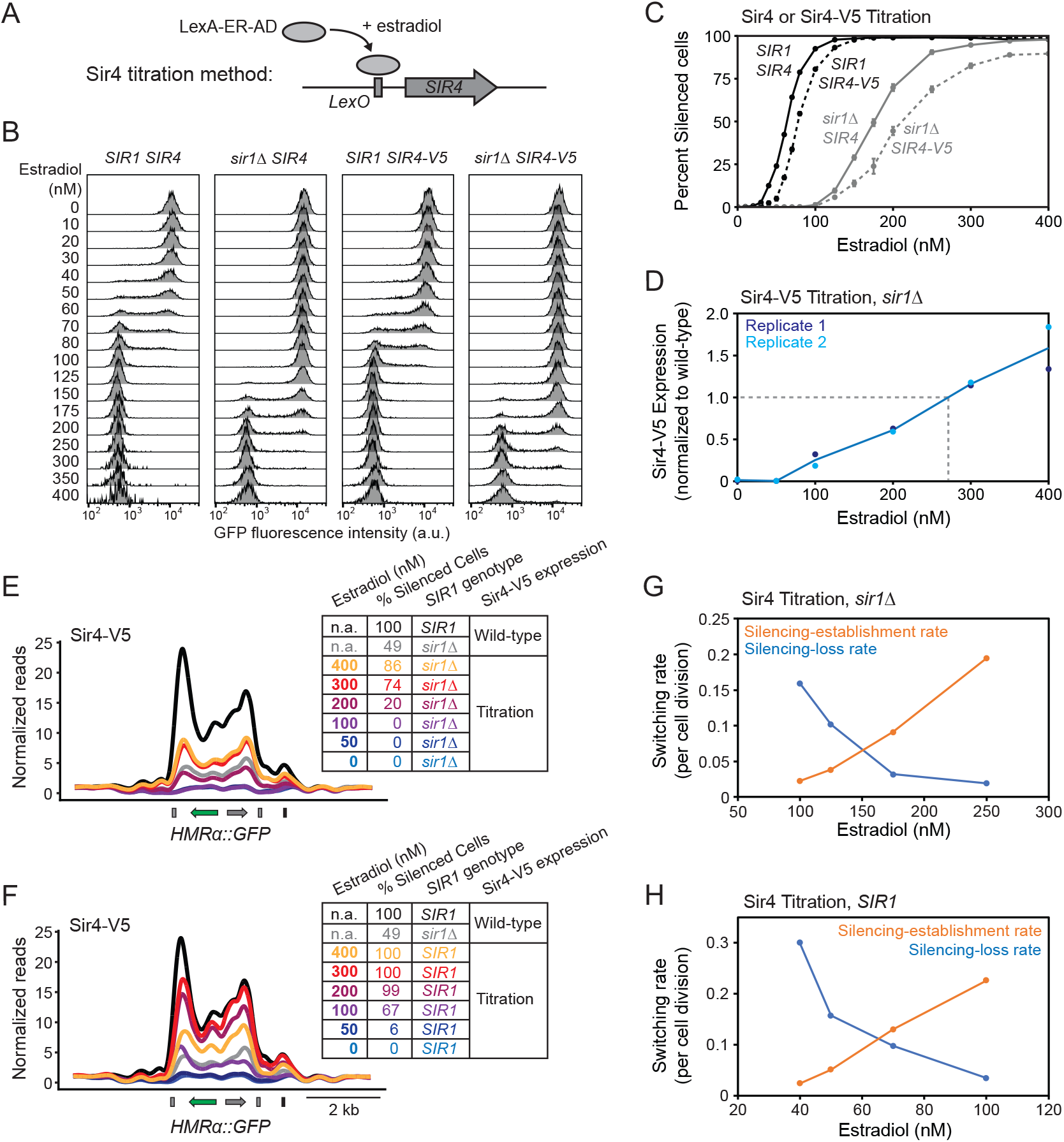
Sir4 dosage determined the frequency of epigenetic states at *HMRα::GFP*. A. Schematic of Sir4 titration via an inducible promoter. B. Fluorescence intensities of strains with *SIR1* or *sir1Δ* and titration of either untagged Sir4 or Sir4-V5 (JRY13092, JRY13093, JRY13637, and JRY13638). Three biological replicates were performed for each condition, and representative flow cytometry profiles are shown. C. Quantification of the percentage of silenced cells for different estradiol concentrations in B. Data are means ± SD (n = 3 independent cultures per condition). D. Sir4-V5 expression at different estradiol concentrations, as measured by western blots (Figure S3B) (JRY13638). Sir4-V5 expression was normalized to Sir4-V5 expression from the native *SIR4* promoter (JRY13648). The blue line represents the mean of both replicates at each estradiol concentration. The grey dashed line represents the projected estradiol concentration (265 nM estradiol) that produced Sir4-V5 at similar levels as wild-type expression. E. ChIP of Sir4-V5 at different estradiol concentrations in *sir1Δ* cells (JRY13638). ChIP of Sir4-V5 expressed from the native *SIR4* promoter was also performed in *SIR1* and *sir1Δ* cells, which are provided as controls (JRY13647 and JRY13648). Reads were normalized to the genome-wide median. A second biological replicate and additional quantification is provided in Figure S3. The percent of silenced cells was calculated by flow cytometry. F. ChIP of Sir4-V5 at different estradiol concentrations in *SIR1* (JRY13637). Controls with wild-type Sir4-V5 expression are identical to E because ChIP in E and F was performed in parallel. A second biological replicate and additional quantification is provided in Figure S3. G-H. Switching rates between epigenetic states at different estradiol concentrations in *sir1Δ* and *SIR1* (JRY13092 and JRY13093). Dividing cells were monitored by live-cell microscopy over a 10-hr time course (n > 750 divisions per genotype). The silencing-establishment rate reflected the frequency of expressed cell divisions in which a daughter cell switched to the silenced state, and the silencing-loss rate reflected the frequency of silenced cell divisions in which a daughter cell switched to the expressed state.

How does the expression level of Sir4 influence silencing in *sir1Δ?* Strikingly, the concentration of estradiol determined the frequency of silenced and expressed cells in *sir1Δ:* higher estradiol concentrations yielded higher frequencies of silenced cells, and lower estradiol concentrations yielded lower frequencies of silenced cells (Figure 2B-C). Additionally, given that *sir1Δ* reduced the efficiency of Sir4 binding and exhibited bistable silencing, we tested whether reducing the expression levels of Sir4 in *SIR1* could phenocopy these effects. Indeed, low concentrations of estradiol generated bistable silencing in *SIR1* and variations in estradiol concentration modulated the frequency of silenced and expressed cells within this bistable regime. Therefore, bistability could be achieved by lowering Sir4 levels or removing a protein that stabilizes Sir4 binding at silenced loci. In both cases, different levels of Sir4 changed the probability that *HMRα::GFP* existed in either the silenced or expressed state.

The range of silencing phenotypes observed at different estradiol concentrations prompted us to quantify the relationship between estradiol concentration and the level of Sir4 expression. We replaced endogenous *SIR4* with *SIR4-V5* to facilitate quantification via western blots. Notably, *SIR4-V5* mildly destabilized silencing and thus decreased the frequency of silenced cells at different estradiol concentrations (Figure 2B and C). The estradiol concentration regime used in our silencing assays ranged from a minimum of undetectable Sir4-V5 to a maximum of 1.5-fold more Sir4-V5 than Sir4-V5 expression from the native *SIR4* promoter (Figure 2D, Figure S3B). We calculated that 265 nM estradiol produced Sir4-V5 at a similar level to the native *SIR4* promoter, and this expression condition yielded ~70% silenced cells in *sir1Δ* (Figure 2C). Additionally, the titration of Sir4-V5 in *SIR1* showed that bistability could be observed when Sir4-V5 was expressed at < 50% of wild-type levels. Therefore, a modest increase in Sir4 dosage rescued *sir1Δ* silencing defects in most cells, and a two-fold reduction in Sir4 dosage in *SIR1* generated bistable silencing.

Our results argued that a bistable population reflects an admixture of silenced cells with reduced Sir4 binding and expressed cells with no Sir4 binding (Figure 1). If this principle applied to the bistable silencing observed at different Sir4 expression levels, the frequency of silenced cells in each population should reflect the level of Sir4 binding measured at the population level. To test this, we titrated Sir4-V5 to different levels in *sir1Δ* cells and performed ChIP on these populations. The highest concentration of estradiol exhibited both a high frequency of silenced cells and intermediate binding of Sir4-V5 at *HMRα::GFP* at the population level (Figure 2E, Figure S3C, D, G). Lower concentrations of estradiol led to fewer silenced cells and proportional decreases in Sir4-V5 binding. We also titrated Sir4-V5 in *SIR1* cells and found robust Sir4-V5 binding at high estradiol concentrations, and decreased Sir4-V5 binding at lower estradiol concentrations (Figure 2F, Figure S3E, F, H). Bistable silencing was observed when Sir4-V5 binding was < 50% of wild-type binding levels, consistent with the results seen in *sir1Δ*. Interestingly, we also observed a decrease in Sir4 binding at the highest estradiol concentration in *SIR1* cells, suggesting that high levels of Sir4-V5 may disrupt important stoichiometries in the Sir complex. Therefore, removal of Sir1 or decreased expression of Sir4 led to decreased binding of Sir4 at *HMRα::GFP* and pushed this locus into a bistable silencing regime. Inside this bistable regime, changes in Sir4 expression changed the frequency of cells in the silenced state, likely by modulating the probability that a destabilized silenced structure could be formed and maintained at *HMRα::GFP*.

Previous studies show that individual *sir1Δ* cells usually maintain and transmit their expression state to daughter cells, but rare switching events do occur (Pillus and Rine, 1989; Xu et al., 2006). We reasoned that different Sir4 expression levels may affect the frequency of these switching events, which could account for different frequencies of silenced cells observed at the population level. To address this possibility, we used live-cell microscopy to monitor the expression states of dividing cells in different estradiol concentrations. In both *SIR1* and *sir1Δ* cells, higher concentrations of estradiol led to lower frequencies of silencing loss and higher frequencies of silencing establishment (Figure 2G-H, Movies S1-S2); these differences in switching rates were consistent with different frequencies of silenced cells at the population level (Table S1). The relationship between Sir4 levels and silencing loss suggested that the probability that a cell maintained and transmitted the silenced state to daughter cells was sensitive to the amount of available Sir4. Additionally, establishing a silenced state was more probable with higher levels of Sir4.

In *sir1Δ* cells, a silenced structure formed at *HMRα::GFP*, but this structure had reduced Sir protein binding and could spontaneously convert to a euchromatic state. We have found that reducing the dosage of Sir4 in *SIR1* generated a similar condition in which Sir4 binding was reduced and silencing became bistable. Within these bistable regimes, the dosage of Sir4 affected the frequency of silenced cells within the population. Higher frequencies of silenced cells at higher Sir4 dosages were correlated with, and presumably due to, fewer silencing-loss events and more silencing-establishment events. A possible cooperativity-based mechanism for these conversion events between epigenetic states is discussed below.

### Silencing was determined in *cis*

If the silenced state at *HMLα::RFP* or *HMRα::GFP* was generated by cooperative binding of Sir complexes and feedback between the Sir complex and histone modifications, the silenced state would be determined in *cis* at the relevant locus. Consistent with this expectation, a previous study found that the epigenetic states of two alleles of *HML* are regulated independently in a *sir1Δ* diploid (Xu et al., 2006). Given that bistability can be observed at *HMRα::GFP* in both *sir1Δ* and *SIR1*, we tested whether bistable states were determined in *cis* in one or both of these contexts. To address this point, we made *HMRα::RFP/HMRα::GFP* diploids that were homozygous for both *LexA-ER-AD* and *LexO-Sir4.* Different concentrations of estradiol revealed that *sir1Δ* diploid cells could silence both alleles, express both alleles, or silence one allele while the other remained expressed (Figure 3A and C, Figure S4). However, a chi-square test on the frequencies of cells in each of these categories argued that these states were not completely independent in *sir1Δ* (*p* < 0.001). Interestingly, a similar analysis of *SIR1* diploids with reduced levels of Sir4 showed very high frequencies of cells that were silenced at both alleles or expressed at both alleles, and very few cells that were silenced only at one but not the other (Figure 3B and D, Figure S4), arguing that the expression states of *HMRα::RFP* and *HMRα::GFP* did not behave independently in *SIR1* (*p* < 0.001). Therefore, epigenetic states in *sir1Δ* were determined by a mix of *cis* and *trans* effects, while epigenetic states in *SIR1* were mostly determined in *trans*.

**Figure 3:**
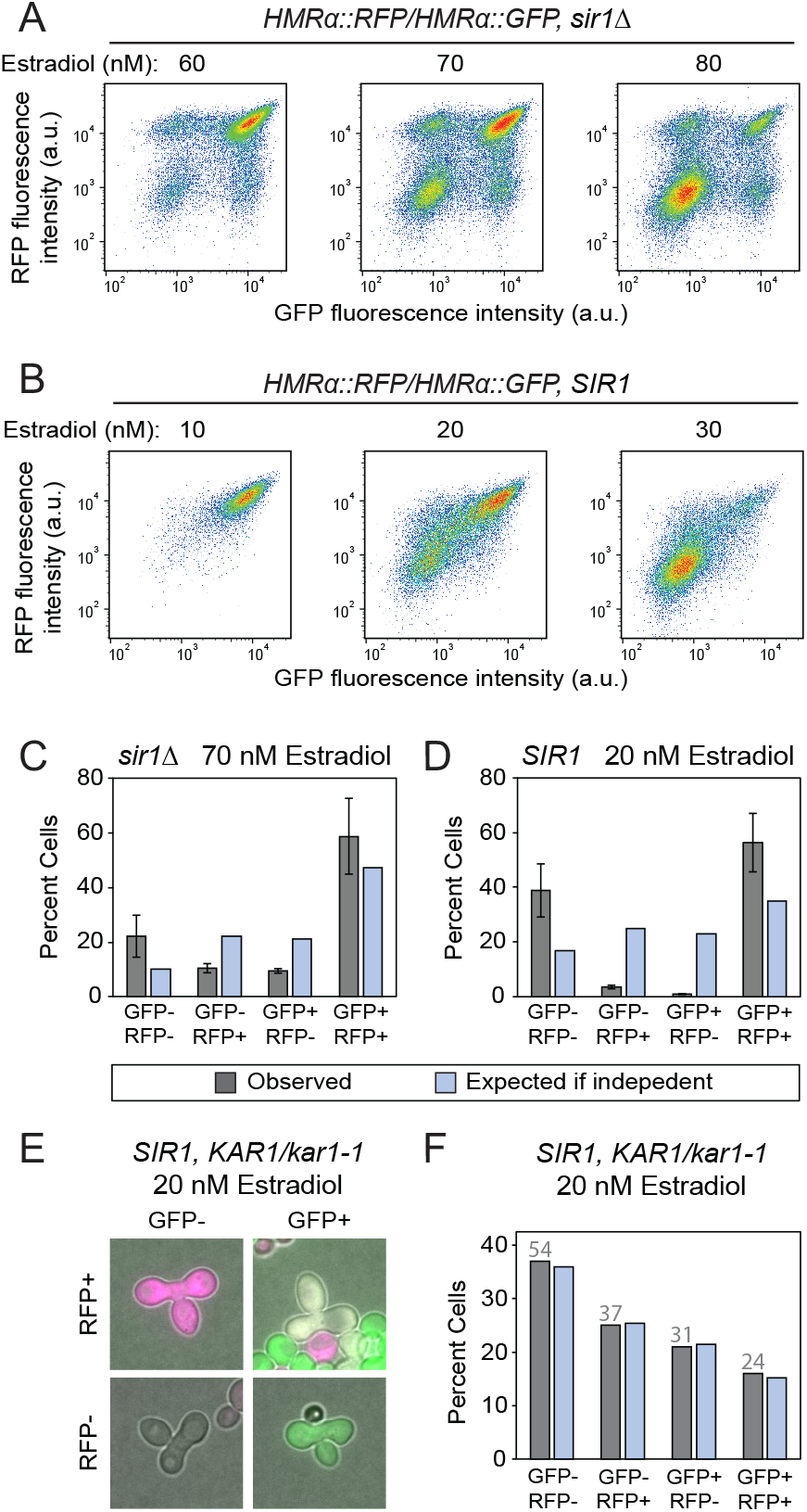
Epigenetic states were determined in *cis* when maintained in separate nuclei. All strains were homozygous for *LexO-SIR4* and *LexA-ER-AD.* A-B. Fluorescence intensity profiles for *HMRα::RFP/HMRα::GFP sir1Δ* diploids and *HMRα::RFP/HMRα::GFP SIR1* diploids (JRY13724 and JRY13725). Three biological replicates were analyzed per condition and representative flow profiles are shown. Additional data with more estradiol concentrations are shown in Figure S4. C. Gray bars represent the percentage of cells in different fluorescence categories for *HMRα::RFP/HMRα::GFP sir1Δ* diploids grown in 70 nM estradiol. Data are means ± SD (n = 3 biological replicates). Light blue bars represent the expected percentage of cells in each category if *HMRα::RFP* and *HMRα::GFP* were regulated independently. These values were calculated by building a 2 x 2 contingency table from the observed data. A chi-square test argued that the states at *HMRα::RFP* and *HMRα::GFP* were not independent (*p* < 0.001). D. Same as C, but performed with *HMRα::RFP/HMRα::GFP SIR1* diploids grown in 20 nM estradiol. A chi-square test argued that the states at *HMRα::RFP* and *HMRα::GFP* were not independent (*p* < 0.001). E. *HMRα::RFP KAR1 SIR1* haploids (JRY13727) were mated to *HMRα::GFP kar1-1 SIR1* haploids (JRY13708) in 20 nM estradiol to generate *HMRα::RFP/HMRα::GFP KAR1/kar1-1 SIR1* heterokaryons. These cells were imaged in 20 nM estradiol over a 10-hr time course. Examples of all four possible state combinations are shown in E, and time-lapse movies of individual heterokaryons are shown in Movie S3. F. Fluorescence profiles of heterokaryons that were > 8 hrs old were quantified by microscopy (n = 146 cells). Gray bars represent observed percentages of cells in each category, and the number of cells counted for each category is placed above the corresponding gray bar. Expected percentages were calculated as described for C and D. A chi-square test failed to reject the null hypothesis that the states at *HMRα::RFP* and *HMRα::GFP* were independent in this context (*p* = 0.72).

One possible explanation for *trans* effects could be cell-to-cell variation in the efficiency of the Sir4 titration method, such as differential responsiveness to estradiol. In this case, cell-to-cell variation in Sir4 expression levels could influence both alleles of *HMRα* equally in *trans*. Alternatively, it is possible that both alleles of *HMRα* could affect each other through a process similar to transvection, in which physical proximity of two alleles allows them to be co-regulated. This possibility would be consistent with observations that silenced loci can physically interact in the nucleus (Hamdani et al., 2019; Miele et al., 2009).

To test whether physical proximity of both *HMRα* alleles contributed to their co-regulation, we created cells in which these alleles were maintained in separate nuclei but shared a common cytoplasm. Diploids that are heterozygous for the *kar1-1* mutation exist as heterokaryons, which are cells that separately maintain nuclei inherited from each original haploid parent (Conde and Fink, 1976). We reasoned that if two alleles of *HMRα* were still co-regulated when maintained in separate nuclei, this would argue against a transvection-like mechanism and suggest that a cytoplasmic mechanism, such as cell-to-cell variation in Sir4 expression levels, was the cause of observed co-regulation. To test this, we made *SIR1 HMRα::RFP/HMRα::GFP KAR1/kar1-1* heterokaryons that were homozygous for the Sir4 titration components and measured silencing at intermediate estradiol concentrations. Over a 10-hour time course, we qualitatively observed many examples in which silencing was maintained at one allele while the other allele remained expressed, and events in which one allele switched to a different state while the other allele retained the same state (Movie S3). To quantify independence, we calculated the frequency of different states in heterokaryons that were formed at least eight hours previously and failed to reject the null hypothesis that epigenetic states at *HMRα::RFP* and *HMRα::GFP* are regulated independently (*p* = 0.72).

Together, these findings showed that the epigenetic states at two *HMRα* alleles were determined in *cis* when these alleles were maintained in separate nuclei that shared a common cytoplasm, and that *trans* effects were observed when these alleles shared a common nucleus. Thus, any *trans* effects observed in *HMRα::RFP/HMRα::GFP KAR1/kar1-1* diploids were not an artifact of the Sir4 titration method, and likely due to proximity-related processes between both *HMRα* alleles. Additionally, our data suggested that this potential proximity effect was stronger in *SIR1* than *sir1Δ;* this idea is consistent with observations that silenced loci physically interact more frequently in *SIR1* than *sir1Δ* (Miele et al., 2009). We conclude that the determinants of epigenetic states at *HMRα* acted in *cis* in both *SIR1* and *sir1Δ* cells, and that *HMRα* alleles in the same nucleus can influence the epigenetic states of each other.

### Silencer-bound proteins modulated Sir4 binding efficiency and the probability of silencing

Our data argued that Sir4 binding efficiency determined the probability of silencing, and that the silencing probability was determined in *cis*. Therefore, we tested whether silencer-binding proteins that recruit the Sir complex in *cis* also modulated the probability of silencing. The *HMR-E* silencer has binding sites for ORC, Rap1, and Abf1, which all interact with Sir proteins (Figure 4A) (Brand et al., 1987; Buchman et al., 1988; Moretti et al., 1994). Specifically, ORC binds to Sir1, Sir1 binds to Sir4, Rap1 binds to Sir4 and Sir3, and Abf1 binds to Sir3. The *HMR-E* silencer is both necessary and sufficient to silence *HMR* (Brand et al., 1985). Previous work has established that individually mutating the ORC, Rap1, or Abf1 binding sites permits silencing of *HMR*, yet mutating any two sites leads to expression of *HMR* as judged by population-level assays (Brand et al., 1987). Additionally, studies have found that some silencer mutations can yield bistable silencing (Mahoney et al., 1991; Sussel et al., 1993). Thus, one possibility is that the ORC, Rap1, and Abf1 binding sites in *HMR-E* increase the efficiency of Sir complex binding, and that combinatorial removal of these sites can reduce Sir complex binding and bring silencing into a bistable regime.

**Figure 4:**
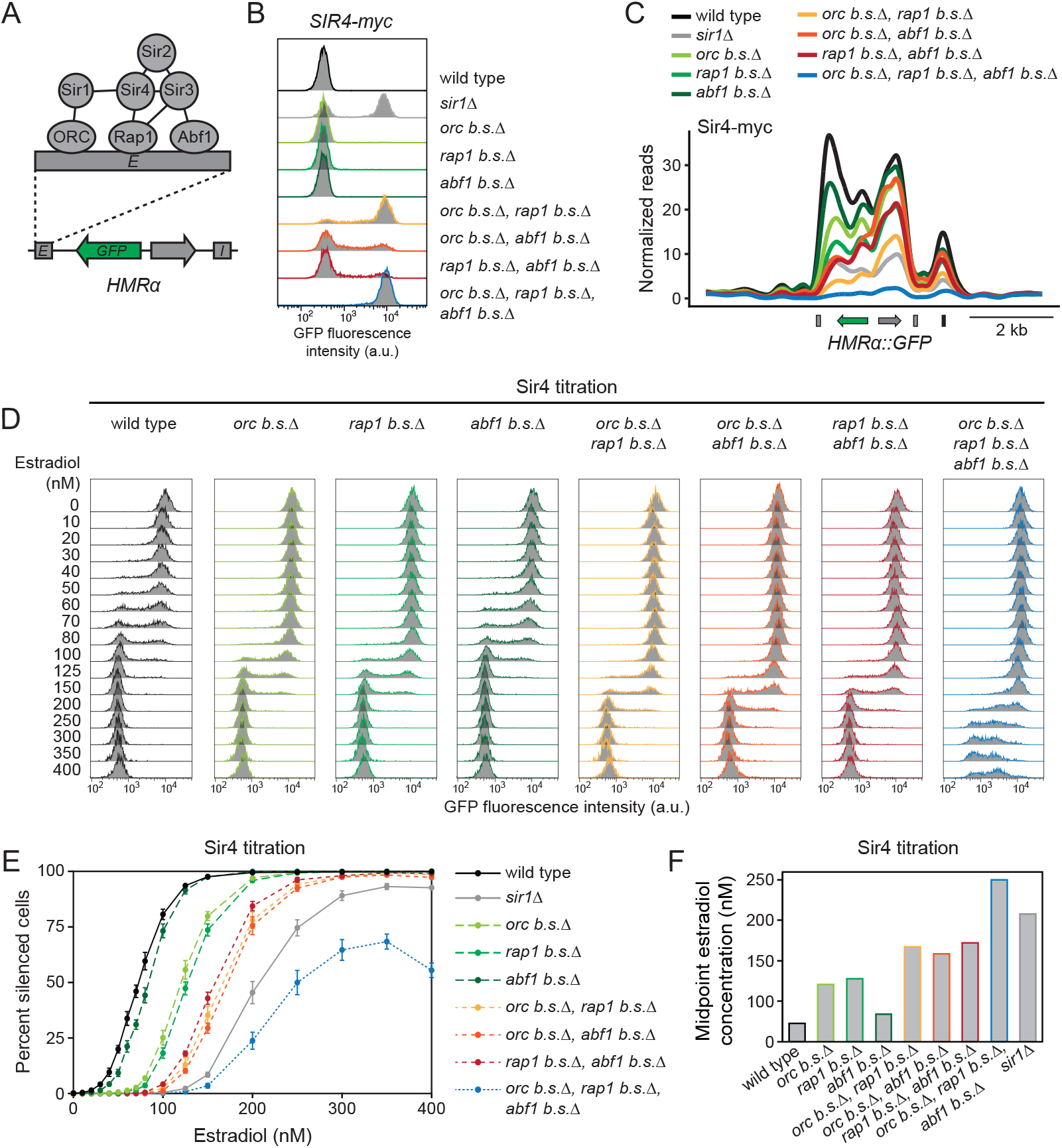
Silencer-binding proteins modulated the frequency of epigenetic states at *HMRα::GFP*. A. Schematic of DNA-binding proteins at the *HMR-E* silencer. Interactions between these DNA-binding proteins and Sir proteins are shown, as well as interactions between Sir proteins. B. Fluorescence intensities of cells with deletions of different binding motifs, and *SIR4-myc* expressed from the native *SIR4* promoter (JRY13130, JRY13132, and JRY13693-13699). Each strain was grown and analyzed in biological triplicate and representative flow cytometry profiles are shown. C. ChIP of Sir4-myc for the same samples shown in B. ChIP-seq reads were normalized to the genome-wide median. A biological replicate and additional quantification are provided in Figure S5. D. Strains with components of the *SIR4* titration method and deletions of different binding motifs at *HMR-E* (JRY13092, JRY13093, and JRY13701-13707). *SIR4* was untagged in these strains. Cells were grown for 24 hrs at log phase in the indicated estradiol concentrations and fixed for flow cytometry. Each condition was tested in biological triplicate and representative flow cytometry profiles are shown. E. Quantification of the percent of silenced cells shown in D. Data are means ± SD (n = 3 biological replicates). F. Projected concentrations of estradiol that correspond to 50% silenced cells, which is termed the midpoint estradiol concentration. These values were calculated from data shown in E.

To test this possibility, we made a series of binding site deletions in the *HMR-E* silencer in a strain containing *SIR4-myc*. Consistent with previous studies, removal of any single binding site still permitted silencing of *HMRα::GFP* in all cells (Figure 4B). Strikingly, removal of any two sites caused *HMRα::GFP* to be silenced in some cells and expressed in others. Of the three strains that lacked two binding sites, the strain lacking the ORC and Rap1 binding sites had lowest number of silenced cells, arguing that these two factors individually played larger roles in silencing than Abf1. Finally, removal of all three binding sites caused nearly all cells to be fully expressed. These data argued that partially functional versions of *HMR-E* can generate bistable silencing, consistent with the bistability observed in *sir1Δ*.

Given our observations that bistability could result from reduced Sir4 binding, we measured Sir4-myc binding in different silencer mutants. Even though removal of any one binding site for ORC, Rap1, or Abf1 still permitted silencing in all cells, these mutants all exhibited reductions in Sir4-myc binding at the *HMR-E* silencer, *α* promoter, and the *HMR-I* silencer (Figure 4C, Figure S5). Removal of any two sites exhibited stronger binding defects, and removal of all three sites led to the strongest binding defects. Therefore, silencer mutants reduced Sir4 binding, and specific mutants with intermediate Sir4 binding exhibited bistability. Additionally, though mutations in the *HMR-E* silencer exhibited Sir4-myc binding defects at *HMR-E* and other regions within *HMRα::GFP*, these defects usually diminished at sites that were further from *HMR-E.* Together, these data argued that Sir4 binding at silencers supports Sir4 binding in other parts of a silenced domain.

Given that removal of any single binding site for ORC, Rap1, or Abf1 led to reduced Sir4 binding but exhibited silencing in all cells, we were curious if altering Sir4 dosage in these mutant backgrounds could reveal silencing defects. Consistent with previous results, titration of untagged Sir4 in different silencer mutant backgrounds generated bistable regimes and determined the frequency of silenced and expressed cells in those regimes (Figure 4D and E). Interestingly, silencer mutants exhibited clear differences in the dose-response relationship between estradiol concentration and the frequency of silenced cells. As a proxy for these shifted dose-response curves, we calculated the estradiol concentration that corresponded to 50% silenced and 50% expressed cells for each genotype and termed this the midpoint estradiol concentration (Figure 4F). Removal of the Abf1 binding site increased this midpoint estradiol concentration only slightly, and removal of the ORC or Rap1 binding sites increased the concentration more significantly. Removal of any two binding sites increased the midpoint estradiol concentration even further than the single binding site mutants, and removal of all three had the highest midpoint estradiol concentration. At high estradiol concentrations, this triple binding-site mutant exhibited cells that were silenced, expressed, or silenced at an intermediate level, a phenomenon that has not been previously documented in *S. cerevisiae.* Therefore, different Sir4 dosages revealed that silencer-binding proteins affected the probability that a given cell was silenced or expressed. By extension, these data argued that silencer-binding proteins changed the concentration of Sir4 that was necessary to achieve silencing.

Here, we found that *sir1Δ* and various mutations in the *HMR-E* silencer affected the efficiency of Sir4 binding and that intermediate degrees of Sir4 binding exhibited bistability. Given that bistable silencing previously correlated with bistable nucleation at silencers, it is likely that weakened versions of the *HMR-E* silencer can also produce bistable nucleation. In this context, our data argued that the number of possible contacts between silencer-bound proteins and the Sir complex determined (1) the probability of Sir4 nucleation at the *HMR-E* silencer, (2) the probability of Sir4 binding across the rest of *HMRα::GFP*, and (3) the probability that *HMRα::GFP* was silenced or expressed. These data further supported the idea that silencers determine the epigenetic state of *HMRα::GFP* in *cis*, as would be expected by a cooperativity model.

### Spreading stabilized Sir4 nucleation at silencers

If Sir complex binding at silencers and nearby nucleosomes were probabilistic events that reinforced each other, mutations that affect spread of Sir proteins across nucleosomes should also affect Sir protein binding at silencers. Spreading of the Sir complex involves a feedback loop between Sir proteins and histone modifications. Sir2 deacetylates various lysine residues on histones H3 and H4, most notably H4K16ac (Imai et al., 2000; Landry et al., 2000). Sir3 binds weakly to acetylated histones via a Bromo Adjacent Homology (BAH) domain and this binding is strengthened approximately 50-fold when histones are deacetylated by Sir2 (Carmen et al., 2002; Onishi et al., 2007). Additionally, deacetylation of H4K16 blocks the ability of Dot1 to methylate H3K79, and unmethylated H3K79 further potentiates Sir3 binding (Altaf et al., 2007; Valencia-Sánchez et al., 2021). The ability of nucleosome-bound Sir3 to recruit additional Sir2 completes the feedback loop, allowing the Sir complex to spread from silencers across silencer-proximal nucleosomes.

Previous studies tested the role of spreading by using mutant versions of Sir2 that are unable to deacetylate histones; these mutants are spreading-defective and show a reduction in Sir complex binding at silencers (Hoppe et al., 2002; Rusché et al., 2002; Thurtle and Rine, 2014). To address this question more systematically, we generated a series of different mutations in Sir proteins and histones that are known to, or predicted to, affect Sir protein spreading. H4K16Q mimics acetylated H4K16. H3K79L and H3K79M both partially mimic H3K79me3 by having a neutralized charge. sir2-N345A protein is expressed at the same levels as wild-type Sir2 but is catalytically inactive and spreading defective (Imai et al., 2000). Similarly, sir3-bahΔ is expressed at similar levels as wild-type Sir3 but is spreading defective (Buchberger et al., 2008). Importantly, given that most of the yeast genome is acetylated at H4K16, methylated at H3K79me3, and lacks Sir protein binding (Ellahi et al., 2015; Leeuwen et al., 2002; Thurtle and Rine, 2014), we reasoned that these mutants were unlikely to generate confounding pleiotropic effects outside of silenced loci.

We first characterized these mutants by measuring silencing of *HMLα::RFP* and *HMRα::GFP* at the single cell level. Interestingly, most spreading mutants exhibited monostable silencing that was neither fully silenced nor fully expressed, but rather expressed at an intermediate level (Figure 5A and B, Figure S6A and B). Within this range of phenotypes, *H3K79L* and *H3K79M* exhibited the weakest silencing defects, and these defects were observed only at *HMLα::RFP. H4K16Q* and *sir3-bahΔ* exhibited stronger silencing defects, and *sir2-N345A* exhibited the strongest silencing defects. In all mutants, the silencing defects were stronger at *HMLα::RFP* than *HMRα::GFP*, consistent with previous findings that silencing at *HMRα* is more robust overall (Saxton and Rine, 2019). Therefore, these spreading mutants provided an array of silencing defects that were observed at both *HMLα::RFP* and *HMRα::GFP*.

**Figure 5:**
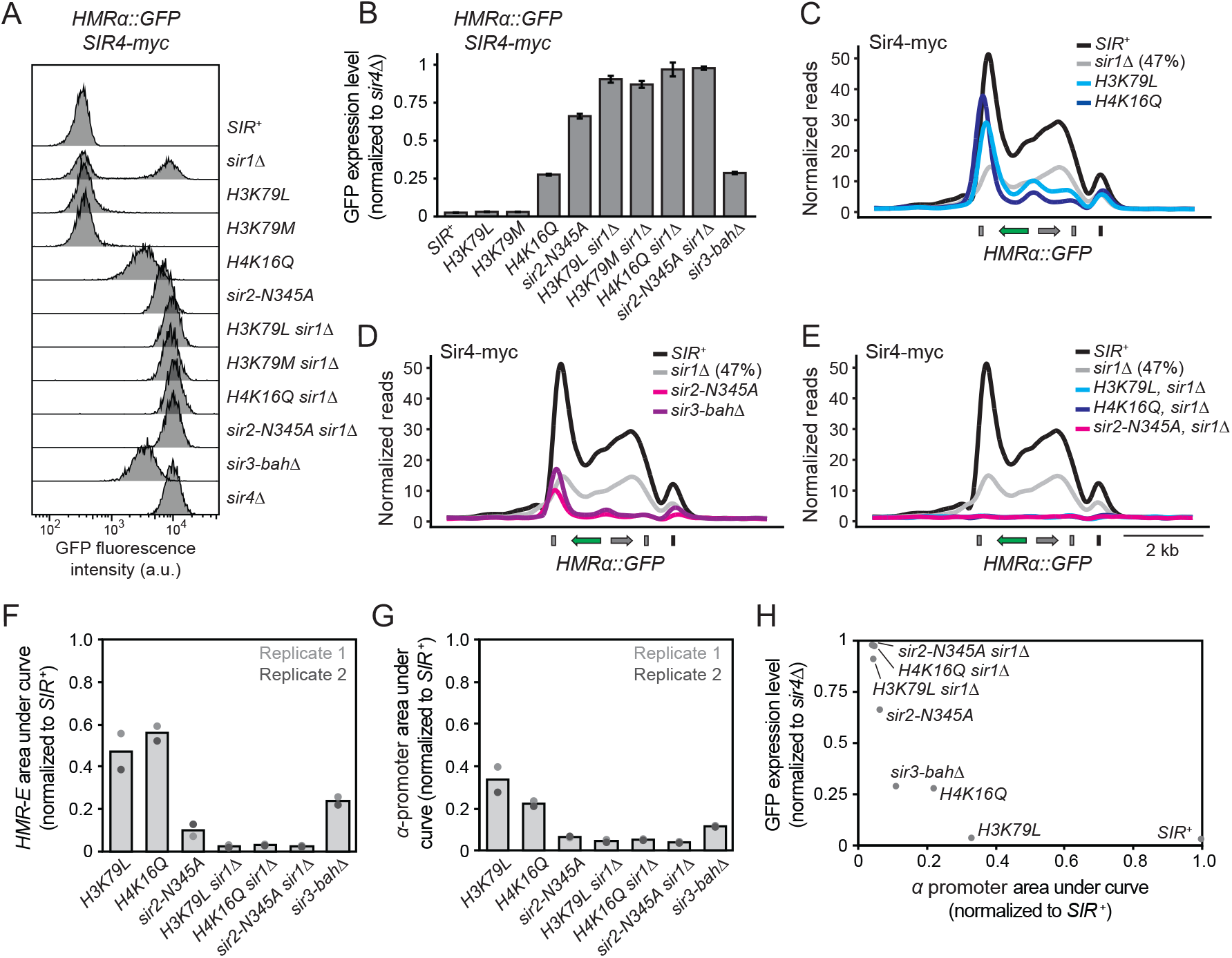
Spreading mutants affected Sir4 nucleation at silencers. A. Fluorescence intensity profiles for different spreading mutants (JRY13233-13239, JRY13631, and JRY13632). *SIR*^+^, *sir1Δ*, and *sir4Δ* strains are provided as controls (JRY11496, JRY13130, and JRY13132). All strains contained *HMRα::GFP* and *SIR4-myc* expressed from the native promoter. Strains were grown and analyzed in biological triplicate, and representative flow cytometry profiles are shown. B. Quantification of GFP expression levels from A, as calculated via the geometric mean intensity statistic in FlowJo. Data are means ± SD (n = 3 biological replicates). C-E. ChIP of Sir4-myc in different spreading mutant backgrounds. Reads were normalized to the genome-wide median. Biological replicates are provided in Figure S7. The percent of silenced cells in *sir1Δ* is shown next to the *sir1Δ* label. F-G. Quantification of Sir4-myc ChIP in spreading mutants. The area under curve was calculated at *HMR-E* (F) and the *α* promoter (G). Data for replicate 1 correspond to C-E, and data for replicate 2 correspond to Figure S7. Gray bars represent the mean of both replicates for each strain. H. Relationship between the GFP expression level (B) and amount of Sir4-myc at the *α* promoter (G).

How do these spreading mutants affect the efficiency of Sir complex spreading and nucleation at silencers? Similar to the range of silencing phenotypes, we observed a range of different binding profiles of Sir4-myc at both *HMLα::RFP* and *HMRα::GFP* (Figure 5C, D, F, G, Figures S6-S7). All spreading mutants exhibited moderate to severe reductions in Sir4 binding across chromatin. *H3K79L* cells exhibited the smallest spreading defects, *H4K16Q* cells had stronger spreading defects, and *sir2-N345A* and *sir3-bahΔ* cells had the strongest spreading defects. Each mutant also exhibited defects in Sir4-myc binding at silencers, and the severity of these defects correlated with the severity of spreading defects. These findings point to an interesting phenomenon in which Sir complex spreading and silencing were not always binary but could also occur at intermediate levels. Within this framework, our data strongly suggested that the efficiency of spreading also impacted the efficiency of nucleation.

If Sir complexes bind cooperatively across silencers and nucleosomes, we would predict that combining *sir1Δ* and spreading mutations would lead to synergistic silencing defects. Alternatively, if Sir complex binding at nucleosomes was entirely downstream of nucleation at silencers, the combination of spreading mutants with *sir1Δ* would yield a mix of cells that fully express *HMRα::GFP* and express *HMRα::GFP* at an intermediate level; similar results would be expected for *HMLα::RFP*. We found that all double mutants tested led to monostable, full expression of both *HMLα::RFP* and *HMRα::GFP* (Figure 5A and B, Figure S6A). Importantly, the finding that *H3K79L* cells exhibited minimal silencing defects alone and *H3K79L sir1Δ* cells were all expressed provided the strongest evidence that silencers and modified histones act synergistically to facilitate silencing. Additionally, spreading mutants combined with *sir1Δ* exhibited no measurable nucleation or spread of Sir4-myc at *HMLα::RFP* and *HMRα::GFP* (Figure 5E-G, Figures S6-S7). These findings argued that Sir complex binding occurs synergistically across silencers and nucleosomes.

In the process of analyzing Sir4-myc binding profiles within different spreading mutants, we noticed an inverse correlation between the amount of Sir4-myc at the *α* promoter and the level of GFP or RFP expression (Figure 5H, Figures S6-S7). More specifically, *H3K79L* reduced Sir4-myc binding efficiency at the *α* promoter yet exhibited no silencing defects at *HMRα::GFP* and minimal silencing defects at *HMLα::RFP.* Other spreading mutants had lower efficiencies of Sir4-myc binding at the *α* promoter and stronger silencing defects. These findings suggested that various spreading mutations that affect silencing may do so by modulating Sir4 binding activity at the *α* promoter.

Together, these findings argued that disrupting different contacts between the Sir complex and nucleosomes modulated the binding efficiency of Sir4 at nucleosomes and silencers. This concept supported a model in which Sir complex binding at nucleosomes and silencers were probabilistic events that reinforced each other. We suspect that these Sir complex binding events also affected the probability that the Sir complex can modify histones, which would further stabilize Sir protein binding and thus contribute to the feedback loop that generates the silenced state.

## Discussion

A common feature of the silenced state of heterochromatin is that it can be generated and inherited at a given locus in some cells, but not others. Previous studies argue that this silenced state is determined by a *cis*-acting mechanism. To identify possible determinants of the silenced state, we assessed the chromatin architecture of the silenced and expressed states of *HMLα* and *HMRα* in *sir1Δ* cells. Strikingly, the results of this experiment argued that the removal of Sir1 weakened silencers and caused Sir protein binding at silencers and nucleosomes to become bistable: intermediate Sir complex binding correlated with the silenced state, and an absence of Sir binding correlated with the expressed state. Additionally, mutations in silencers, mutations affecting Sir complex spreading, and changes in Sir4 dosage led to coordinated changes in the efficiency of Sir complex nucleation and spread in most cases. These findings supported a recent theory suggesting that Sir complex binding could be bistable, and that this property would be driven by (1) cooperative binding of the Sir complex and (2) feedback between the Sir complex and histone modifications (Sneppen and Dodd, 2015). Our data strongly argued that the silenced state is determined by synergy between these two processes, explaining previously paradoxical observations that silenced epigenetic states require silencers continuously.

### Silencers generated bistability

Historically, studies of heterochromatin regard silencers as structures that are responsible only for the initial nucleation of silencing proteins. Then, spreading of silencing proteins and modification of histones would act downstream of nucleation and be sufficient to transmit the memory of silencing to daughter cells. In contrast, we found that silencing protein nucleation and spread did not behave as hierarchical steps but acted synergistically in a manner that resembled a feedback loop. As such, mutations that affected nucleation also affected spread, and mutations that affected spread also affected nucleation. These findings argued that nucleation and spread are not separate processes with separate roles in establishment and inheritance of silencing. Instead, Sir complexes appeared to bind cooperatively across silencers and nucleosomes, and silencing dynamics likely reflected the dynamics of this coordinated Sir complex binding activity.

Building on past studies (Mahoney et al., 1991; Pillus and Rine, 1989; Sussel et al., 1993), we found that various silencer mutations and reductions in Sir4 dosage could convert constitutive silencing into bistable silencing. The Sir4 binding profiles in these different conditions pointed to a common theme: strong silencers promoted robust Sir complex binding across silencers and nucleosomes, and weaker silencers reduced this Sir complex binding efficiency. Similarly, reductions in Sir4 dosage also led to decreases of Sir4 binding efficiency. Within this framework, when Sir4 binding was sufficiently reduced, the bistability of silencing emerged: some cells were silenced at *HMLα* and *HMRα* and other cells were expressed at these loci. These results argued that Sir complex binding can occur at multiple intermediate levels, and the reduction of binding below a critical threshold reveals bistable properties of silenced loci.

When Sir complex binding is reduced and generates a bistable silencing regime, what are the molecular architectures of silenced and expressed chromatin states? By focusing on one bistable condition, *sir1Δ*, we found that the silenced states of *HMLα* and *HMRα* exhibited strong reductions in Sir complex binding at silencers, mild reductions in Sir complex binding across nucleosomes within these loci, and no reductions in histone modifications associated with silencing. In contrast, the expressed states of these loci showed no measurable binding of the Sir complex at silencers or nucleosomes, and these nucleosomes lacked silencing-specific histone modifications. Thus, even though silencing exhibited all-or-nothing properties, the silenced state in *sir1Δ* was characterized by intermediate Sir4 binding. We inferred that this intermediate binding was still strong enough to promote cooperative binding and maintain a feedback loop involving histone modifications, yielding silencing that was indistinguishable from *SIR1* cells. However, this form of feedback is likely prone to failure in *sir1Δ*, such that the silenced state can probabilistically convert to an expressed state (Pillus and Rine, 1989; Xu et al., 2006) in which components of the silencing feedback loop are absent entirely.

Within a bistable silencing regime, changes in silencer strength and Sir4 dosage were able to titrate the frequency of silenced cells and the population-level Sir4 binding efficiency. Given our observations that silenced states in *sir1Δ* cells exhibited intermediate Sir complex binding and expressed states in *sir1Δ* cells exhibited no Sir complex binding, we suspected that different ratios of cells with these molecular architectures produced different Sir4 binding levels as measured by population-level ChIP assays. Indeed, the percentage of silenced cells in a population was positively correlated with the Sir4 binding efficiency. Thus, our data indicated that within a bistable regime, factors that changed the Sir complex binding efficiency at the population level did so by affecting the probability that a locus exhibited either intermediate Sir complex binding or no binding at all. In turn, this probability would determine the frequency of silenced cells in a population.

Epigenetic states in *sir1Δ* cells exist in a dynamic equilibrium: a silenced state has a low probability of switching to an expressed state (silencing loss), and an expressed state has a low probability of switching to the silenced state (silencing establishment) (Pillus and Rine, 1989; Xu et al., 2006). We found that higher dosages of Sir4 decreased the probability of silencing loss and increased the probability of silencing establishment in both *sir1Δ* and *SIR1* cells, leading to more silenced cells overall. One possible explanation for this trend is that the loss of a silenced structure results from stochastic reductions in Sir complex binding below levels needed for cooperative behavior. Conversely, the formation of a silenced structure may reflect a stochastic association of Sir complexes at levels that are sufficient to initiate cooperative behavior. In this context, high Sir4 dosages could push *HMLα* and *HMRα* into the silenced state in all cells by effectively eliminating the possibility of silencing loss and ensuring that any expressed state rapidly establishes silencing. Additionally, given that weaker silencers produce bistable silencing and stronger silencers produce constitutive silencing, it is likely that silencer strength also modulates the probability of silencing loss and silencing establishment. This model offers a mechanistic explanation for the emergence of bistability in silencer mutants and, by extension, the roles of silencers in regulating epigenetic states.

### Feedback mechanisms in silencing

Previous studies of biological circuits demonstrate that bistability requires at least one form of a feedback loop (Ferrell, 2002). Our data supported a model in which bistable silencing is driven by two forms of feedback: cooperative binding of the Sir complex and a feedback loop between the Sir complex and histone modifications. More specifically, Sir complex binding at silencers and adjacent nucleosomes would be cooperative, probabilistic events that occur infrequently in the expressed state of a locus. However, these events could probabilistically generate silencing-specific histone modifications, which would increase the probability of Sir complex binding at nucleosomes. Stronger binding of the Sir complex at modified nucleosomes could then promote binding at silencers through cooperativity, and thus complete a two-part feedback loop that involves silencers and histone modifications. Additionally, a recent study demonstrates that PRC2-based silencing can be antagonized by a transcription-based feedback loop that promotes a heritable expressed state (Holoch et al., 2021). Currently, it is unclear whether the expressed state of *HML* and *HMR* in *sir1Δ* cells is similarly driven by a transcription-based feedback loop, or is simply a default condition when a silencing-based feedback loop is inactive.

How can the binding of a Sir complex at one location stabilize the binding of another Sir complex nearby? Multiple homotypic and heterotypic interactions exist between components of the Sir complex: Sir2, Sir3, and Sir4 can all physically interact with each other, Sir3 can homodimerize, and Sir4 can homodimerize (Moazed and Johnson, 1996; Moazed et al., 1997; Moretti et al., 1994). In principle, this collection of mutual interactions could lead to cooperative binding of the Sir complex across silencers and nucleosome arrays. Interestingly, *in vitro* studies argue that the Sir complex binds cooperatively but only across 2-3 nucleosomes, and that binding of more Sir proteins to larger nucleosome arrays does not improve cooperativity further (Behrouzi et al., 2016; Swygert et al., 2018). These data may explain why mutations in the *HMR-E* silencer exhibited Sir4 binding defects that diminished at larger distances from *HMR-E.* Additionally, reducing the size of a silenced locus may destabilize silencing by reducing the valency of binding sites for the Sir complex; however, reducing the size of *HMRα* from twelve to six nucleosomes does not affect the probability of silencing loss in *sir1Δ* (Saxton and Rine, 2019). These findings argue that the Sir complex binds cooperatively across nucleosomes, but caution that this process only occurs over limited distances.

Numerous studies in different organisms have identified feedback loops in which silencing proteins can both modify histones and bind to the modifications that they generate (Hansen et al., 2008; Hecht et al., 1995; Imai et al., 2000; Zhang et al., 2008). In budding yeast, Sir2 deacetylates histones at multiple residues and Sir3 binds more strongly to deacetylated histones (Carmen et al., 2002; Imai et al., 2000; Landry et al., 2000). Additionally, both activities antagonize the H3K79 methylase Dot1, and demethylated H3K79 further strengthens Sir3 binding (Altaf et al., 2007; Valencia-Sánchez et al., 2021). We observed that different mutations in this pathway reduced Sir4 binding efficiency to different extents. Surprisingly, most of these mutations led to monostable silencing that was not fully silenced or fully expressed, but silenced to some intermediate degree. Why did spreading mutations exhibit monostable, intermediate silencing while silencer mutations produced bistability? One possibility is that the probability of silencing is driven primarily by events initiated at the silencer, yet the success of these events requires cooperative binding of the Sir complex and successful modification of histones. In contrast, the strength of silencing *per se* would be driven by the efficiency of Sir complex binding throughout the locus, which varied significantly in different spreading mutants.

### The transmission of silencing to daughter cells

Our data supported a model in which cooperative binding and a feedback loop between silencing proteins and histone modifications could generate a silenced state in some cells but not others. Could these processes also support the transmission of the silenced state to daughter cells? Multiple studies show that the induced removal of silencers permits maintenance of silencing in arrested cells but leads to rapid silencing loss in dividing cells (Audergon et al., 2015; Coleman and Struhl, 2017; Holmes and Broach, 1996; Laprell et al., 2017; Ragunathan et al., 2015). Additionally, studies in *S. pombe* show that a weak silencer that is incapable of *de novo* silencing establishment can support the inheritance of a silenced state (Wang and Moazed, 2017; Wang et al., 2021). These data argue that during DNA replication, silencers facilitate the transmission of the silenced state to daughter chromatids. In light of our data suggesting that the Sir complex can bind cooperatively across silencers and nucleosomes, it is possible that Sir complexes either physically bridge across the replication fork or that Sir complexes evicted by replication rapidly re-bind to daughter chromatids in a manner that is potentiated by other bound Sir complexes. Consistent with this idea, previous work argues that some silencing proteins can be transmitted to daughter chromatids during DNA replication, though this has yet to be tested for the Sir complex (Francis et al., 2009; Mazo et al., 2012).

Conversely, what is the role of histone inheritance in the inheritance of the silenced state? Mutations in replication-coupled histone chaperones Dpb3 and Mcm2 lead to strong defects in histone inheritance and minor effects on the inheritance of silencing in *S. cerevisiae* (Saxton and Rine, 2019; Schlissel and Rine, 2019). In *S. pombe*, localized methylation of H3K9me3 can silence a reporter gene in the absence of a silencer, and this silenced state is heritable in cells that contain the H3K9 methylase Clr4 and lack the H3K9 demethylase Epe1 (Audergon et al., 2015; Ragunathan et al., 2015). These studies argue that histone inheritance contributes to silencing inheritance under normal conditions, and that this contribution can be improved significantly by removing a silencing antagonist. Based on these findings, we suspect that the inheritance of modified histones acts synergistically with cooperative binding of silencing proteins to drive silencing inheritance; as such, stronger or weaker versions of these two processes could tune the efficiency of silencing inheritance.

By monitoring epigenetic states in dividing cells, we observed that the transmission of the silenced state to daughter cells was more probable at higher Sir4 dosages. One possible implication of this result is that higher concentrations of Sir4 increase the probability that Sir proteins successfully transfer from the parental chromatid to daughter chromatids, and thus increase the probability of silencing inheritance. In this context, the inheritance of modified histones and cooperative binding properties of the Sir complex may act synergistically to enhance the likelihood that a daughter chromatid receives a sufficient quantity of Sir proteins to maintain silencing. Conversely, rare events of silencing loss may reflect events in which a daughter chromatid inherits an insufficient number of Sir proteins to maintain cooperative binding and histone modification behavior. Therefore, this cooperativity model offers an explanation for the continuous requirement of silencers in epigenetic inheritance.

## Materials and Methods

### Yeast strains

Strains and oligonucleotides used in this study are listed in Table S2. All strains were derived from the W303 background. Components of the Sir4 titration method were engineered as previously described (Ottoz et al., 2014). Deletions and point mutations were made with CRISPR/Cas9 technology (Lee et al., 2015). Each deletion or mutation utilized a single guide RNA (sgRNA) and a repair template, which are listed in Table S2. Repair templates that were ≤ 80 nucleotides were generated by PCR-amplifying two oligonucleotides with partial homology. Larger oligonucleotides were synthesized or amplified from plasmids. Co-transformation of the repair template and a plasmid containing Cas9 and an sgRNA resulted in transformants with the desired mutation. All mutations were confirmed by junction PCR and sequencing.

All strains used in this study contained either *HMLα* or *HMRα* (that contained fluorescent reporter genes), but never both. Given that these two loci share a significant amount of homology, they were separated in different strains to avoid ambiguity in mapping ChIP-seq reads. Additionally, all cells were *matΔ* to avoid ambiguity in ChIP-seq read mapping between *MAT* and *HMLα* or *HMRα. SIR3* and *SIR4* were tagged with either 13x *myc* (referred to as *SIR3-myc* or *SIR4-myc* throughout this study) or 1x V5 (*SIR4-V5*). To generate heterokaryons analyzed in Figure 3, cells that mated similarly to *MATα* cells and cells that mated similarly to *MATα* cells were needed. Therefore, JRY13727 was engineered with two stop codons early in the *HMRα1* gene such that *α1* protein would not be produced; this effectively made these cells mate as *MATα* cells regardless of the expression state from *HMRα*. Additionally, JRY13708 contained the plasmid pJR157 (*MATα, URA3*) to provide constant *α2* protein expression, allowing these cells to mate as *MATα* cells.

To generate combinatorial *HMR-E* silencer mutants, *HMR-E* was replaced with *K. lac LEU2* by auxotrophic selection, and repair templates with different deletions in *HMR-E* were used to replace *HMR-EΔ::K. lac LEU2* with CRISPR/Cas9. To generate point mutations in histones, a single sgRNA was used that could target both histone genes for H3 (*hht1* and *hht2*) or for H4 (*hhf1* and *hhf2*). Similarly, a single repair template per mutation was engineered to have homology to both genes for a given histone. Thus, transformation of Cas9, sgRNA, and the repair template yielded colonies that mutated both genes simultaneously. Mutations were confirmed by sequencing with histone gene-specific primers.

### Flow cytometry

With the exception of sorting experiments in Figure 1 and Figure S1-S2, all flow cytometry was performed on cells that had been grown at log-phase for 24 hrs. Specifically, three independent cultures per condition were inoculated in Yeast extract Peptone + 2% Dextrose (YPD) liquid media and grown to saturation overnight at 30°C. These cultures were then serially back-diluted in YPD liquid media and grown at log-phase at 30°C for 24 hrs to allow cells to reach equilibrium. For experiments that utilized estradiol, the overnight growth to saturation and log-phase growth were both performed in YPD + the indicated estradiol concentration. All growth was performed in 96-well plates (Corning CLS3788) on an incubating microplate shaker (VWR 12620-930).

After 24 hrs of log-phase growth, cells were centrifuged, resuspended in 100 μl of 4% paraformaldehyde, and fixed at room temperature for 15 min. Fixed cells were subsequently pelleted and resuspended in 150 μl 1X PBS solution. These samples were stored at 4°C until they were analyzed by flow cytometry. Flow cytometry was performed with an LSR Fortessa (BD Biosciences) with a FITC filter for GFP and a PE-TexasRed filter for RFP. At least 20,000 cells were analyzed per genotype, with the exception of some samples that had slower growth rates due to growth in high estradiol concentrations (i.e. 400 nM estradiol in Figure 2B). FlowJo software (BD Biosciences) was utilized to analyze flow cytometry data. All cells within an experiment were gated identically.

### Live cell imaging

Given that YPD media exhibits high autofluorescence, live-cell imaging experiments were performed with Complete Supplement Mixture + 2% dextrose (CSM) media. Cells were first grown as described above (overnight to saturation, and for 24 hrs at log-phase) in CSM liquid media with or without estradiol. 3 μl of cells at a concentration of 0.5 OD were subsequently spotted onto CSM media plates that contained the same estradiol concentration as the liquid media. Once dry, a sterile spatula was used to cut a 5 mm x 5 mm square surrounding the cell patch. This agar square was removed and inverted onto a 35 mm glass bottom dish (Thermo Scientific 150682). Cells were subsequently imaged using a Zeiss Z1 inverted fluorescence microscope with a Prime 95B sCMOS camera (Teledyne Photometrics), Plan-Apochromat 63x/1.40 oil immersion objective (Zeiss), filters, MS-2000 XYZ automated stage (Applied Scientific Instrumentation), and MicroManager imaging software (Open Imaging). Since cells were held between the agar and glass slide, a single Z-frame was used rather than a Z-stack.

For time-lapse imaging, samples were maintained at 30°C and humidified by a P-set 2000 Heated Incubation Insert (PeCon). Brightfield and fluorescence images for 16 different fields of view per sample were collected every 20 mins for 10 hrs. Images were analyzed with ImageJ (NIH). Counting of fluorescence states and switching events between states was performed manually with a single-blind approach. If the fluorescence profile of a cell was not clearly silenced or expressed, or there was ambiguity regarding a potential switching event, that cell or cell division was excluded from analysis.

To generate and image heterokaryons, *HMRα::RFP KAR1 SIR1* (JRY13727) haploids were mated to *HMRα::GFP kar1-1 SIR1* haploids (JRY13727) by mixing on CSM + 20 nM estradiol plates for 4 hrs at 30°C. This population of cells was then resuspended in sterilized water, diluted to 0.5 OD, and spotted onto a separate part of the same agar plate. Cells were imaged as described above. Given that the mating efficiency was low, relatively few heterokaryons were observed per field of view; therefore, 80 fields of view were imaged. Imaging was performed every 20 min over a 10-hr time course. Heterokaryons that had formed prior to imaging or within the first 100 mins were manually tracked for at least 8 hrs, after which their fluorescence profiles were analyzed for Figure 3F. Separately, a small sample of these cells were fixed and stained with DAPI. This staining confirmed that heterokaryons had two nuclei.

### Western blots

Protein isolation and western blotting were performed as described previously (Brothers and Rine, 2019). Membranes were blocked with Odyssey Blocking Buffer (LI-COR Biosciences). Primary antibodies were rabbit anti-Hxk2 (Rockland #100-4159, 1:10,000), mouse anti-myc (Thermo Fisher MA1-980, 1:500), and mouse anti-V5 (Thermo Fisher R960-25, 1:1000). Secondary antibodies used were goat anti-rabbit CW680 (LI-COR #C81106-03, 1:20,000) and goat anti-mouse CW800 (LI-COR #C60531-05, 1:20,000). Western blots were imaged with a LI-COR infrared fluorescent scanner.

### Cell sorting

Prior to sorting, a single culture per genotype was inoculated into YPD liquid media and grown to saturation overnight at 30°C. Cultures were then back-diluted and grown at log-phase for > 24 hrs at 30°C. *sir1Δ* cells at ~1 O.D. were sorted using a FACSAria Fusion cell sorter (BD Biosciences) equipped with a FITC filter for GFP and a PE-TexasRed filter for RFP. Fluorescence gates were calibrated from *SIR^+^* and *sir4Δ* cells grown in parallel. Approximately 3,000,000 *sir1Δ* cells that were silenced at the locus of interest (*HMLα::RFP* or *HMRα::GFP*) were sorted into one population, and 3,000,000 *sir1Δ* cells that were expressed at that locus were sorted into a different population. Control strains were subjected to the same growth regimen, were not sorted, and were diluted to the same O.D. concentration as the *sir1Δ* sorted populations. Each population was split into two populations to provide biological replicates, and these populations were grown in 50 mL YPD liquid media for approximately 12 hrs at 30°C, or until they reached approximately 1 O.D. At this point, these cells were fixed for flow cytometry and ChIP-seq, as described below.

### ChIP-seq

For ChIP experiments that were not part of cell sorting experiments, two independent cultures per condition (the two biological replicates) were inoculated and grown to saturation overnight at 30°C, with or without estradiol. These samples were then back-diluted into 50 mL YPD liquid media, with or without estradiol, and grown at log phase for 24 hrs at 30°C, or until they reached approximately 1 O.D. These samples were then fixed for flow cytometry and ChIP-seq.

For a given sample, cells were always harvested at ~1 O.D. in 50 mL YPD. If cultures were not at the same O.D., the more concentrated cultures were back-diluted such that all samples were harvested simultaneously at ~1 O.D. in 50 mL YPD. For each sample, 150 μl was fixed in 4% paraformaldehyde, as described above, and saved for later analysis by flow cytometry. The remaining sample (~50 mL) was separately treated with formaldehyde (added to a final concentration of 1%) and fixed for 15 min at 30°C while shaking. Fixation was quenched by adding glycine to a final concentration of 300 mM and allowing cells to continue shaking for 10 min. Cells were washed once in 1X PBS, washed once in 1X TE, and cell pellets were flash frozen in liquid nitrogen. Samples were stored at −80°C prior to ChIP processing.

To perform ChIP, cells were first thawed on ice and resuspended in 1 mL FA buffer (50 mM HEPES, 150 mM NaCl, 1 mM EDTA, 1% Triton X-100, 0.1% SDS, 0.1% sodium deoxycholate, with Roche cOmplete protease inhibitors (Sigma)). ~200 μL of 0.5 mm zirconia/Silica beads (BioSpec Products NC0450473) were added to each sample, and cells were lysed using a FastPrep-24 5G (MP Biomedicals) with 6.0 m/s beating for 40 s followed by 2 min on ice, repeated four times total. The cell lysate was transferred to 15 mL Bioruptor Pico tubes along with ~200μL of the corresponding sonication beads (Diagenode C010200031) and sonicated using a Bioruptor Pico (Diagenode B01060010) for 10 cycles of 30 s ON followed by 30 s OFF. After sonication, samples were spun at 4°C for 15 min at 17 k RCF to pellet cellular debris, and ~700 μL of the chromatin-containing supernatant was saved.

Immunoprecipitation was performed by adding BSA to a final concentration of 0.5 mg/mL in the 700 μL of chromatin sample, and adding either 5 μL of undiluted anti-myc antibody (Thermo Fisher MA1-980), 5 μL of undiluted anti-V5 antibody (Thermo Fisher R960-25), 5 μl of undiluted anti-H4 antibody (Sigma 05-858), 5 μl of undiluted anti-H4K16ac antibody (Sigma 07-329), or 5 μl of undiluted anti-H3K79me3 antibody (Diagenode C15410068). Samples were rotated overnight at 4°C. The following day, 50 μL of Protein A Dynabeads (Invitrogen 10002D) per sample was washed twice in FA buffer and resuspended in the same volume of FA buffer. 50 μL of this Dynabead solution was added to each chromatin sample and these samples were rotated at 4°C for 2 hrs. Beads were subsequently washed with the following buffers, and samples were rotated at room temperature for 5 min for each wash step: 1X in FA buffer, 1X in FA buffer + 500 mM NaCl (yielding 650 mM NaCl total), 1X in LiCl buffer (0.25 M LiCl, 0.5% Igepal, 0.5% sodium deoxycholate, 1 mM EDTA, 10 mM Tris-HCl pH 8), and 1X in TE. Samples were eluted by resuspending beads in 100 μL of 1X TE + 1% SDS and incubating at 65°C for 2 hrs with gentle shaking. Then, samples were incubated overnight at 65°C to reverse crosslinking. Finally, 3 μL of 1 mg/mL RNase A (Sigma R6148) and 3 μL of 20 mg/mL Proteinase K (NEB EO0491) was added to each sample, samples were incubated at 42°C for 2 hrs, and DNA was purified by using SPRI beads.

IP samples were prepared for high-throughput sequencing by using the Ultra II DNA Library Prep kit (NEB E7645L). Samples were multiplexed and paired-end sequencing was performed using a MiniSeq (Illumina) or by the Vincent J. Coates Genomics Sequencing Laboratory at UC Berkeley, which used a NovaSeq 6000 (Illumina). Sequencing reads were aligned using Bowtie2 (Langmead and Salzberg, 2012) and reference genomes derived from the *Saccharomyces cerevisiae* S288C genome (GenBank accession number GCA_000146045.2). Coverage was calculated and normalized to the non-heterochromatic genome-wide median, which corresponds to the genome-wide median that excludes rDNA, subtelomeric regions, and all of chromosome III. This coverage calculation and normalization was performed using custom Python scripts (Goodnight and Rine, 2020) and data were plotted using R. Within R, the previously normalized coverage was smoothed with a 300 bp rolling mean. To calculate the area under curve for a given region, the area_under_curve function was used in R (MLmetrics package). Given that the ChIP-seq peaks for Sir3 and Sir4 at silencers were often centered slightly inside of the silencers, the area under curve for *HML-E* and *HMR-E* included an additional 150 bp to the right of each silencer.

## Data availability

Sequencing data have been deposited into the publicly available NCBI Gene Expression Omnibus (GSE195880). All study data are included in the article and supplemental information.

## Acknowledgements

We thank Susan Strome, Martin Howard, Caroline Dean, and Djem Kissiov for useful conversations that contributed to this work. Additionally, we thank the Rine lab for providing valuable input on the experiments presented here. We also thank the UC Berkeley Flow Cytometry facility, especially Hector Nolla and Alma Valeros, for assistance with FACS. Finally, we thank Marc Fouet for extensive assistance with microscopy. This work was supported by an NSF predoctoral fellowship (DGE1752814 to D.S.S) and an NIH grant (GM139488 to J.R.).

## Author Contributions

Daniel S Saxton, Conceptualization, Resources, Data curation, Software, Formal analysis, Validation, Investigation, Visualization, Methodology, Writing - original draft, Writing - review and editing, Project administration, Funding acquisition. Jasper Rine, Conceptualization, Resources, Supervision, Funding acquisition, Writing - review and editing.

## Notes

### Competing Interest Statement

The authors have declared no competing interest.

## References

Altaf, M., Utley, R.T., Lacoste, N., Tan, S., Briggs, S.D., and Côté, J. (2007). Interplay of Chromatin Modifiers on a Short Basic Patch of Histone H4 Tail Defines the Boundary of Telomeric Heterochromatin. Mol Cell 28, 1002–1014.

Audergon, P.N.C.B., Catania, S., Kagansky, A., Tong, P., Shukla, M., Pidoux, A.L., and Allshire, R.C. (2015). Epigenetics. Restricted epigenetic inheritance of H3K9 methylation. Science 348, 132–135.

Behrouzi, R., Lu, C., Currie, M.A., Jih, G., Iglesias, N., and Moazed, D. (2016). Heterochromatin assembly by interrupted Sir3 bridges across neighboring nucleosomes. ELife.

Berry, S., Hartley, M., Olsson, T., Dean, C., and Howard, M. (2015). Local chromatin environment of a Polycomb target gene instructs its own epigenetic inheritance. ELife.

Brand, A.H., Breeden, L., Abraham, J., Sternglanz, R., and Nasmyth, K. (1985). Characterization of a “silencer” in yeast: A DNA sequence with properties opposite to those of a transcriptional enhancer. Cell 41, 41–48.

Brand, A.H., Micklem, G., and Nasmyth, K. (1987). A yeast silencer contains sequences that can promote autonomous plasmid replication and transcriptional activation. Cell 51, 709–719.

Brothers, M., and Rine, J. (2019). Mutations in the PCNA DNA Polymerase Clamp of Saccharomyces cerevisiaeReveal Complexities of the Cell Cycle and Ploidy on Heterochromatin Assembly. Genetics 213, 449–463.

Buchberger, J.R., Onishi, M., Li, G., Seebacher, J., Rudner, A.D., Gygi, S.P., and Moazed, D. (2008). Sir3-nucleosome interactions in spreading of silent chromatin in Saccharomyces cerevisiae. Mol Cell Biol 28, 6903–6918.

Buchman, A.R., Kimmerly, W.J., and Rine, J. (1988). Two DNA-binding factors recognize specific sequences at silencers, upstream activating sequences, autonomously replicating sequences, and telomeres in.… Molecular and Cellular.…

Carmen, A.A., Milne, L., and Grunstein, M. (2002). Acetylation of the Yeast Histone H4 N Terminus Regulates Its Binding to Heterochromatin Protein SIR3. Journal of Biological Chemistry 277, 4778–4781.

Cheng, T.H., and Gartenberg, M.R. (2000). Yeast heterochromatin is a dynamic structure that requires silencers continuously. Genes & Development 14, 452–463.

Coleman, R.T., and Struhl, G. (2017). Causal role for inheritance of H3K27me3 in maintaining the OFF state of a DrosophilaHOX gene. Science eaai8236–20.

Conde, J., and Fink, G.R. (1976). A mutant of Saccharomyces cerevisiae defective for nuclear fusion. Proc National Acad Sci 73, 3651–3655.

Dhillon, N., and Kamakaka, R.T. (2000). A Histone Variant, Htz1p, and a Sir1p-like Protein, Esc2p, Mediate Silencing at HMR. Mol Cell 6, 769–780.

Dodson, A.E., and Rine, J. (2015). Heritable capture of heterochromatin dynamics in Saccharomyces cerevisiae. ELife 4, e05007.

Ellahi, A., Thurtle, D.M., and Rine, J. (2015). The chromatin and transcriptional landscape of native Saccharomyces cerevisiae telomeres and subtelomeric domains. Genetics.

Ferrell, J.E. (2002). Self-perpetuating states in signal transduction: positive feedback, double-negative feedback and bistability. Current Opinion in Cell Biology 14, 140–148.

Francis, N.J., Follmer, N.E., Simon, M.D., Aghia, G., and Butler, J.D. (2009). Polycomb proteins remain bound to chromatin and DNA during DNA replication in vitro. Cell 137, 110–122.

Gardner, K.A., Rine, J., and Fox, C.A. (1999). A region of the Sir1 protein dedicated to recognition of a silencer and required for interaction with the Orc1 protein in saccharomyces cerevisiae. Genetics 151, 31–44.

Goodnight, D., and Rine, J. (2020). S-phase-independent silencing establishment in Saccharomyces cerevisiae. Elife 9, e58910.

Grewal, S., and Klar, A. (1996). Chromosomal inheritance of epigenetic states in fission yeast during mitosis and meiosis. Cell.

Hamdani, O., Dhillon, N., Hsieh, T.-H.S., Fujita, T., Ocampo, J., Kirkland, J.G., Lawrimore, J., Kobayashi, T.J., Friedman, B., Fulton, D., et al. (2019). Transfer RNA Genes Affect Chromosome Structure and Function via Local Effects. Molecular and Cellular Biology 1–67.

Hansen, K.H., Bracken, A.P., Pasini, D., Dietrich, N., Gehani, S.S., Monrad, A., Rappsilber, J., Lerdrup, M., and Helin, K. (2008). A model for transmission of the H3K27me3 epigenetic mark. Nature Cell Biology 10, 1291–1300.

Hecht, A., Laroche, T., Strahl-Bolsinger, S., and Gasser, S.M. (1995). Histone H3 and H4 N-termini interact with SIR3 and SIR4 proteins: a molecular model for the formation of heterochromatin in yeast. Cell 80.

Holmes, S.G., and Broach, J.R. (1996). Silencers are required for inheritance of the repressed state in yeast. Gene Dev 10, 1021–1032.

Holoch, D., Wassef, M., Lövkvist, C., Zielinski, D., Aflaki, S., Lombard, B., Héry, T., Loew, D., Howard, M., and Margueron, R. (2021). A cis-acting mechanism mediates transcriptional memory at Polycomb target genes in mammals. Nat Genet 1–12.

Hoppe, G.J., Tanny, J.C., Rudner, A.D., Gerber, S.A., Danaie, S., Gygi, S.P., and Moazed, D. (2002). Steps in Assembly of Silent Chromatin in Yeast: Sir3-Independent Binding of a Sir2/Sir4 Complex to Silencers and Role for Sir2-Dependent Deacetylation. Molecular and Cellular Biology 22, 4167–4180.

Imai, S., Armstrong, C.M., Kaeberlein, M., and Guarente, L. (2000). Transcriptional silencing and longevity protein Sir2 is an NAD-dependent histone deacetylase. Nature 403, 795–800.

Jackson, V. (1988). Deposition of newly synthesized histones: hybrid nucleosomes are not tandemly arranged on daughter DNA strands. Biochemistry 27, 2109–2120.

Landry, J., Sutton, A., Tafrov, S.T., Heller, R.C., Stebbins, J., Pillus, L., and Sternglanz, R. (2000). The silencing protein SIR2 and its homologs are NAD-dependent protein deacetylases. Proc National Acad Sci 97, 5807–5811.

Langmead, B., and Salzberg, S.L. (2012). Fast gapped-read alignment with Bowtie 2. Nature Methods 9, 357–359.

Laprell, F., Finkl, K., and Müller, J. (2017). Propagation of Polycomb-repressed chromatin requires sequence-specific recruitment to DNA. Science eaai8266–7.

Lee, M.E., DeLoache, W.C., Cervantes, B., and Dueber, J.E. (2015). A Highly Characterized Yeast Toolkit for Modular, Multipart Assembly. ACS Synthetic Biology 4, 975–986.

Leeuwen, F. van, Gafken, P.R., and Gottschling, D.E. (2002). Dot1p modulates silencing in yeast by methylation of the nucleosome core. Cell 109, 745–756.

Mahoney, D.J., Marquardt, R., Shei, G.J., Rose, A.B., and Broach, J.R. (1991). Mutations in the HML E silencer of Saccharomyces cerevisiae yield metastable inheritance of transcriptional repression. Gene Dev 5, 605–615.

Mazo, S.P.Y.S.D.J.J.H.K.B.S.K.S.B.E.C.H.B.A., Sedkov, Y., Johnston, D.M., Hodgson, J.W., Black, K.L., Kovermann, S.K., Beck, S., Canaani, E., Brock, H.W., and Mazo, A. (2012). TrxG and PcG Proteins but Not Methylated Histones Remain Associated with DNA through Replication. Cell 150, 922–933.

Miele, A., Bystricky, K., and Dekker, J. (2009). Yeast silent mating type loci form heterochromatic clusters through silencer protein-dependent long-range interactions. Plos Genet 5, e1000478.

Moazed, D., and Johnson, A.D. (1996). A Deubiquitinating Enzyme Interacts with SIR4 and Regulates Silencing in S. cerevisiae. Cell 86, 667–677.

Moazed, D., Kistler, A., Axelrod, A., Rine, J., and Johnson, A.D. (1997). Silent information regulator protein complexes in Saccharomyces cerevisiae: A SIR2/SIR4 complex and evidence for a regulatory domain in SIR4 that inhibits its interaction with SIR3. Proc National Acad Sci 94, 2186–2191.

Moretti, P., Freeman, K., Coodly, L., and Shore, D. (1994). Evidence that a complex of SIR proteins interacts with the silencer and telomere-binding protein RAP1. Gene Dev 8, 2257–2269.

Ng, H.H., Ciccone, D.N., Morshead, K.B., Oettinger, M.A., and Struhl, K. (2003). Lysine-79 of histone H3 is hypomethylated at silenced loci in yeast and mammalian cells: A potential mechanism for position-effect variegation. Proc National Acad Sci 100, 1820–1825.

Onishi, M., Liou, G.-G., Buchberger, J.R., Walz, T., and Moazed, D. (2007). Role of the Conserved Sir3-BAH Domain in Nucleosome Binding and Silent Chromatin Assembly. Mol Cell 28, 1015–1028.

Ottoz, D.S.M., Rudolf, F., and Stelling, J. (2014). Inducible, tightly regulated and growth condition-independent transcription factor in Saccharomyces cerevisiae. Nucleic Acids Research 42, e130–e130.

Pillus, L., and Rine, J. (1989). Epigenetic inheritance of transcriptional states in S. cerevisiae. Cell 59, 637–647.

Prior, C.P., Cantor, C.R., Johnson, E.M., and Allfrey, V.G. (1980). Incorporation of exogenous pyrene-labeled histone into Physarum chromatin: a system for studying changes in nucleosomes assembled in vivo. Cell 20, 597–608.

Ragunathan, K., Jih, G., and Moazed, D. (2015). Epigenetics. Epigenetic inheritance uncoupled from sequence-specific recruitment. Science 348, 1258699.

Rine, J., and Herskowitz, I. (1987). Four genes responsible for a position effect on expression from HML and HMR in Saccharomyces cerevisiae. Genetics 116, 9–22.

Rusché, L.N., Kirchmaier, A.L., and Rine, J. (2002). Ordered nucleation and spreading of silenced chromatin in Saccharomyces cerevisiae. Molecular Biology of the Cell 13, 2207–2222.

Saxton, D.S., and Rine, J. (2019). Epigenetic memory independent of symmetric histone inheritance. ELife 8, 585.

Schlissel, G., and Rine, J. (2019). The nucleosome core particle remembers its position through DNA replication and RNA transcription. Proceedings of the National Academy of Sciences of the United States of America 12, 201911943–201911947.

Siliciano, P.G., and Tatchell, K. (1984). Transcription and regulatory signals at the mating type locus in yeast. Cell 37, 969–978.

Sneppen, K., and Dodd, I.B. (2015). Cooperative stabilization of the SIR complex provides robust epigenetic memory in a model of SIR silencing in Saccharomyces cerevisiae. Epigenetics 10, 293–302.

Sussel, L., Vannier, D., and Shore, D. (1993). Epigenetic switching of transcriptional states: cis-and trans-acting factors affecting establishment of silencing at the HMR locus in Saccharomyces cerevisiae. Molecular and Cellular Biology 13, 3919–3928.

Swygert, S.G., Senapati, S., Bolukbasi, M.F., Wolfe, S.A., Lindsay, S., and Peterson, C.L. (2018). SIR proteins create compact heterochromatin fibers. Proc National Acad Sci 115, 201810647.

Thurtle, D.M., and Rine, J. (2014). The molecular topography of silenced chromatin in Saccharomyces cerevisiae. Gene Dev 28, 245–258.

Triolo, T., and Sternglanz, R. (1996). Role of interactions between the origin recognition complex and SIR1 in transcriptional silencing. Nature.

Valencia-Sánchez, M.I., Ioannes, P.D., Wang, M., Truong, D.M., Lee, R., Armache, J.-P., Boeke, J.D., and Armache, K.-J. (2021). Regulation of the Dot1 histone H3K79 methyltransferase by histone H4K16 acetylation. Science 371, eabc6663.

Vos, D.D., Frederiks, F., Terweij, M., Welsem, T. van, Verzijlbergen, K.F., Iachina, E., Graaf, E.L. de, Altelaar, A.F.M., Oudgenoeg, G., Heck, A.J.R., et al. (2011). Progressive methylation of ageing histones by Dot1 functions as a timer. Embo Rep 12, 956–962.

Wang, X., and Moazed, D. (2017). DNA sequence-dependent epigenetic inheritance of gene silencing and histone H3K9 methylation. Science eaaj2114–8.

Wang, X., Paulo, J.A., Li, X., Zhou, H., Yu, J., Gygi, S.P., and Moazed, D. (2021). A composite DNA element that functions as a maintainer required for epigenetic inheritance of heterochromatin. Mol Cell 81, 3979–3991.e4.

Xu, E.Y., Zawadzki, K.A., and Broach, J.R. (2006). Single-Cell Observations Reveal Intermediate Transcriptional Silencing States. Molecular Cell 23, 219–229.

Zhang, K., Mosch, K., Fischle, W., and Grewal, S.I.S. (2008). Roles of the Clr4 methyltransferase complex in nucleation, spreading and maintenance of heterochromatin. Nature Structural & Molecular Biology 15, 381–388.

