## Supplemental information for "Distinct silencer states determine epigenetic states of heterochromatin"

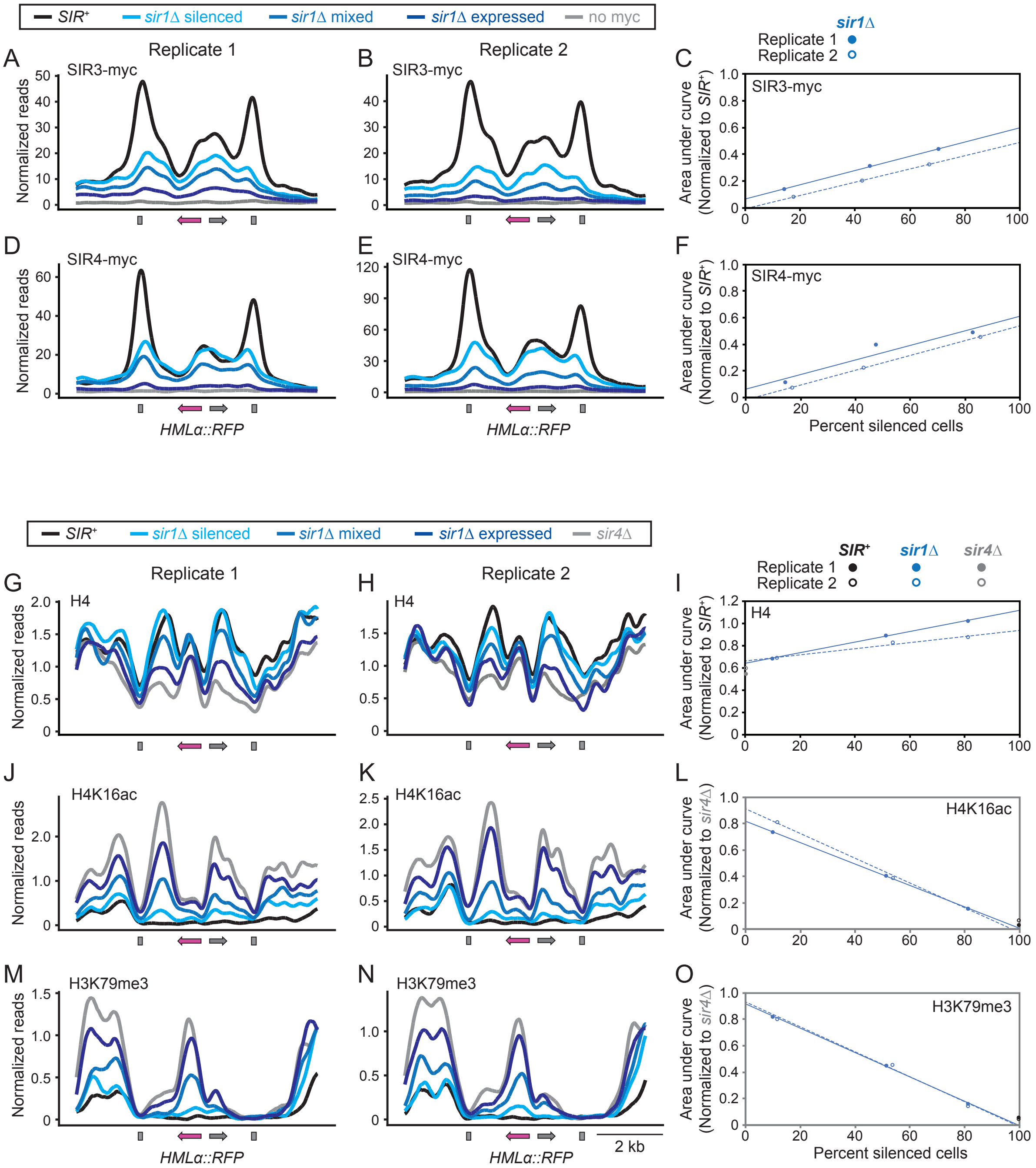


**Figure S1: Biological replicates for ChIP at *HMLα::RFP*.** Data in A, E, K, and N are repeated from Figure 1C, E, G, and I, respectively, and are provided here for reference. However, additional conditions were tested and biological replicates were performed (JRY11472, JRY11494, JRY13125, JRY13127, JRY13129, and JRY13131). First, in addition to the *sir1Δ* silenced and *sir1Δ* expressed populations, a *sir1Δ* mixed population was generated by mixing equal fractions of *sir1Δ* silenced and expressed cells immediately after sorting. Additionally, for A-F, a *SIR*+ strain that lacked a *myc* tag was used as a control (JRY11472). Similar to Figure 1, anti-myc antibodies were used to assess Sir3-myc (A-C) or Sir4-myc (D-F) binding at *HMLα::RFP*. For G-O, chromatin isolated from different populations was split and treated with either anti-H4 antibodies (G-I), anti-H4K16ac antibodies (J-L), or anti-H3K79me3 antibodies (M-O). H4 ChIP was provided as a control to account for the possibility that differences in H4K16ac and H3K79me3 between different *SIR* genotypes were due to differences in nucleosome occupancy. The correlation between Sir complex occupancy and nucleosome occupancy may reflect higher nucleosome turnover in transcriptionally active chromatin. Regardless, H4 ChIP was inversely correlated with H4K16ac and H3K79me3, arguing that nucleosome occupancy did not account for differences in the histone modification landscape between different *SIR* genotypes. Quantification of ChIP was performed by calculating the integrated area under the *HML-E* silencer (C, F) or under the area between the *HML-E* silencer and the $\alpha$ promoter (I, L, O) and normalizing these values to the *SIR*$^{+}$ (C, F, I) or *sir4Δ* (L, O) values. The percent of silenced cells for each population was calculated by flow cytometry. Best fit lines are provided for the *sir1Δ* ChIP data, and the projected value of these lines at 0% silenced cells and 100% silenced cells offers a prediction of the ChIP values in pure populations containing only a single epigenetic state.

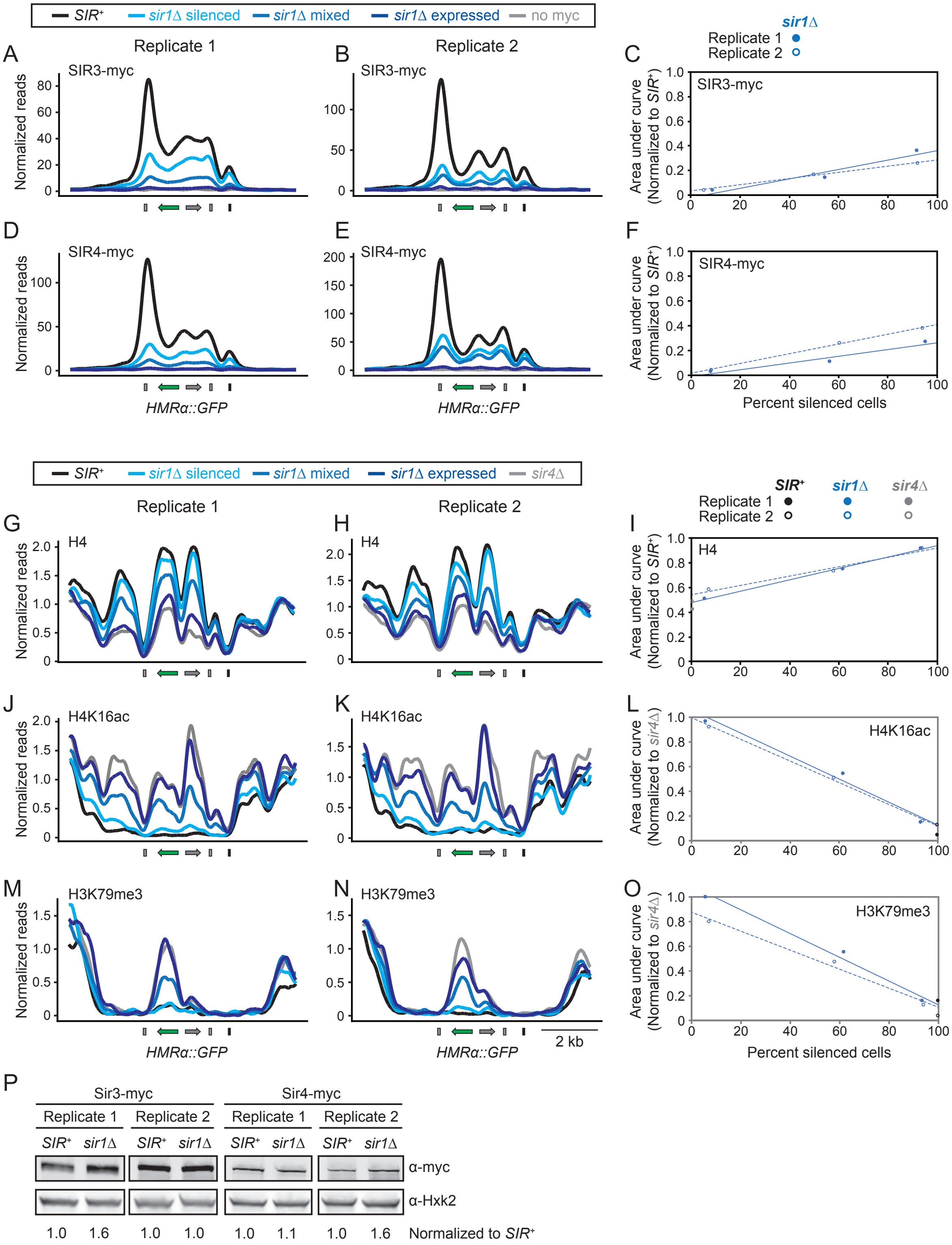


**Figure S2: Biological replicates for ChIP at *HMRα::GFP*.** Data in A, E, K, and M are repeated from data seen in Figure 1D, F, H, and J, respectively, and are provided here for reference. However, additional conditions were tested and biological replicates were performed (JRY11474, JRY11496, JRY13126, JRY13128, JRY13130, and JRY13132). First, in addition to the *sir1Δ* silenced and *sir1Δ* expressed populations, a *sir1Δ* mixed population was generated by mixing equal fractions of *sir1Δ* silenced and expressed cells immediately after sorting. Additionally, for A-F, a *SIR*$^+$ strain that lacked a *myc* tag was used as a control (JRY11474). Similar to Figure 1, anti-myc antibodies were used to assess Sir3-myc (A-C) or Sir4-myc (D-F) binding at *HMR*$\alpha$*::GFP*. For G-O, chromatin isolated from different populations was split and treated with either anti-H4 antibodies (G-I), anti-H4K16ac antibodies (J-L), or anti-H3K79me3 antibodies (M-O). Quantification of ChIP was performed by calculating the integrated area under the *HMR-E* silencer (C, F) or under the area between the *HMR-E* silencer and the $\alpha$ promoter (I, L, O) and normalizing these values to the *SIR*$^+$ (C, F, I) or *sir4Δ* (L, O), values. The percent of silenced cells for each population was calculated by flow cytometry. Best fit lines are provided for the *sir1Δ* ChIP data, and the projected value of these lines at 0% silenced cells and 100% silenced cells offers a prediction of the ChIP values in pure populations containing only a single epigenetic state. P. Two biological replicates of Sir3-myc and Sir4-myc western blots (JRY13126, JRY13128, JRY13130, and JRY13132). Hxk2 expression was used as a control. Sir3-myc or Sir4-myc levels were normalized to Hxk2, and these calibrated levels were compared between *SIR*$^+$ and *sir1Δ*.

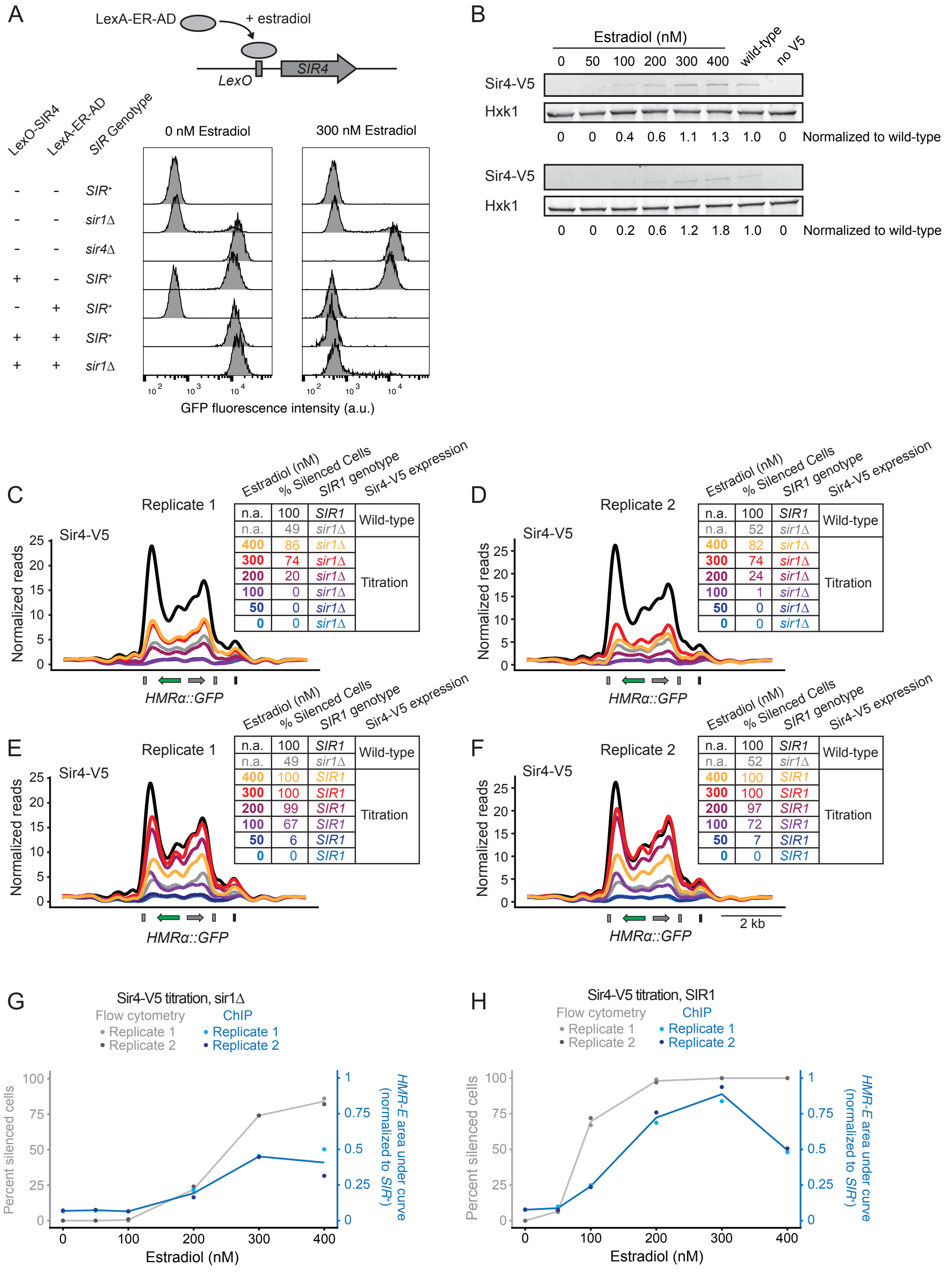


**Figure S3: Sir4 expression driven by an inducible promoter.** A. Cells were grown at log-phase for 24 hrs with or without addition of estradiol and fixed for flow cytometry (JRY11474, JRY11478, JRY11496, JRY13087, JRY13092, JRY13093, JRY13099). Three biological replicates for each condition were performed, and representative profiles are shown. B. Sir4-V5 expression at different estradiol concentrations (JRY13638). Two biological replicates are shown. Sir4-V5 levels were normalized to the Hxk2 controls, and these values were normalized to wild-type Sir4-V5 expression that was driven by the native *SIR4* promoter (JRY13648). A strain with untagged *SIR4* was also tested as a control (JRY11474). This quantification is graphed in Figure 2D. C-D. Two biological replicates for Sir4-V5 ChIP from *sir1Δ* cells, with additional controls (JRY13638, JRY13647, and JRY13648). C is identical to Figure 2E and is provided here for reference. E-F. Two biological replicates for Sir4-V5 ChIP from *SIR1* cells, with additional controls (JRY13637, JRY13647, and JRY13648). E is identical to Figure 2F. G-H. ChIP signal was quantified by calculating the integrated area under the *HMR-E* silencer and normalizing this value to the value observed with wild-type Sir4-V5 expression in *SIR1* cells. The percent of silenced cells for each replicate, which are provided in the legends of C-F, are also plotted for reference. The grey line represents the mean percentage of silenced cells at each estradiol concentration, and the blue line represents the mean area under curve.

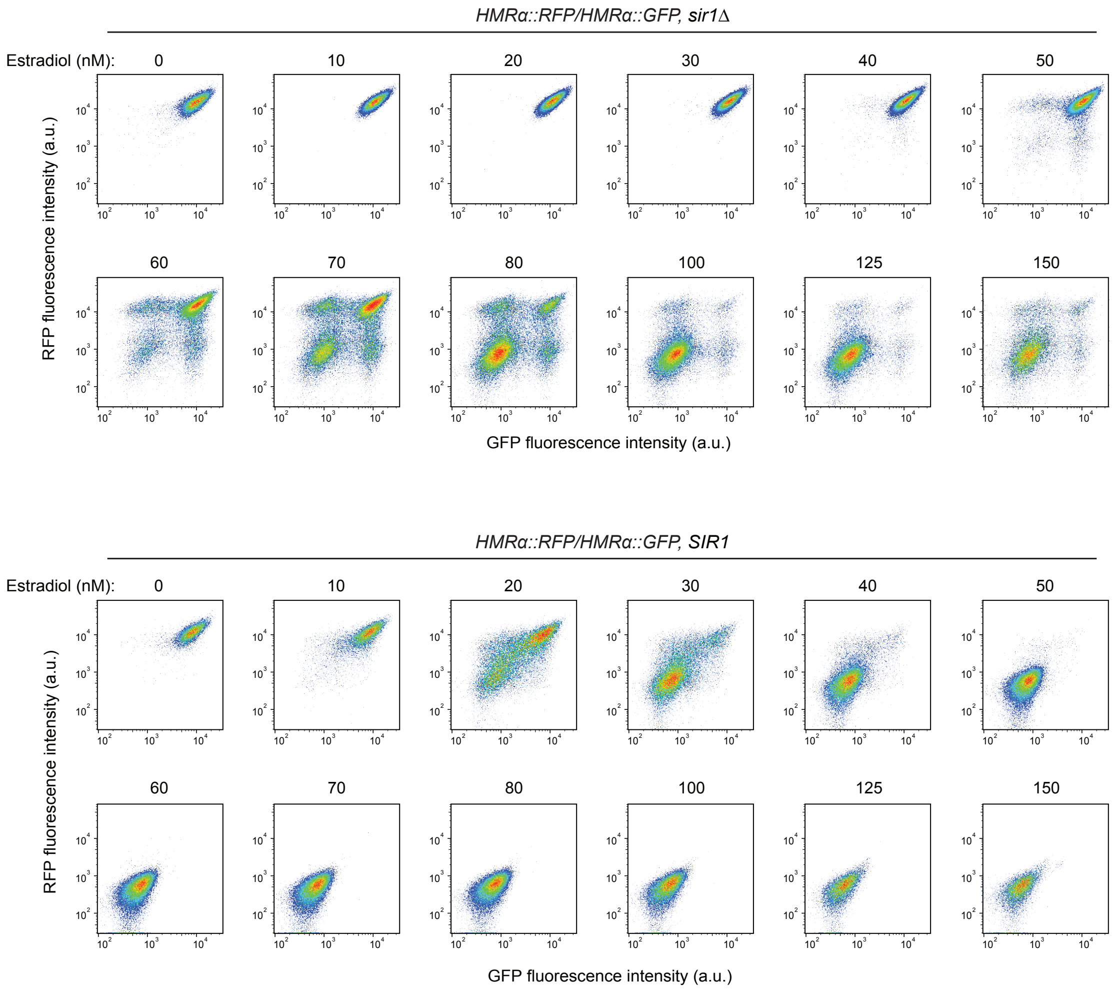


**Figure S4: Fluorescence profiles for *HMRα::RFP/HMRα::GFP* diploids grown in different estradiol concentrations** (JRY13724 and JRY13725)**.** Three biological replicates were tested per condition, and representative flow cytometry profiles are shown. These data are an expanded version of data shown in Figure 3A-B.

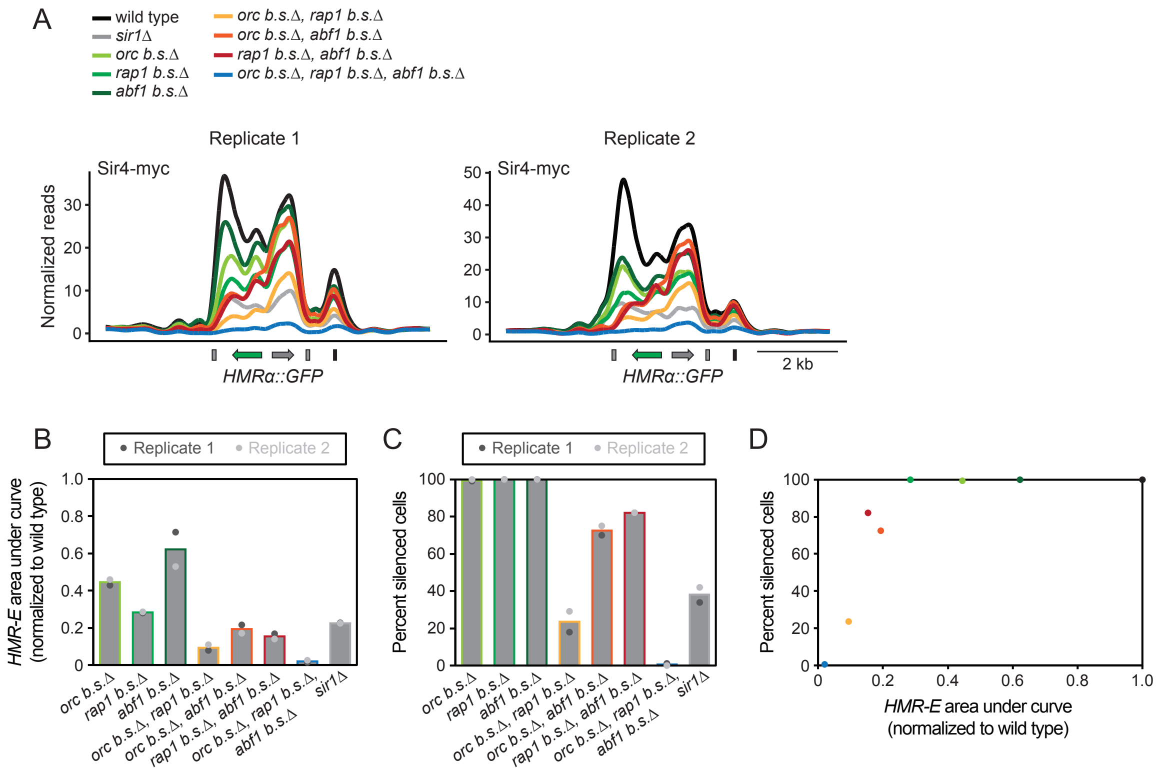


**Figure S5: Biological replicates for Sir4-myc ChIP in silencer mutants.** A. Two biological replicates of Sir4-myc ChIP in *HMR-E* mutants and control strains (JRY13130, JRY13132, and JRY13693-13699). Replicate 1 is identical to data shown in Figure 4C and is shown here for reference. B. ChIP signal was quantified by calculating the integrated area under the *HMR-E* silencer and normalizing this value to the wild-type value. Bars reflect the mean of both replicates for each condition. C. The percent of silenced cells for each replicate, as calculated by flow cytometry. Bars reflect the mean of both replicates. D. Relationship between the average Sir4-myc ChIP signal (B) and percent of silenced cells (C) for each genotype.

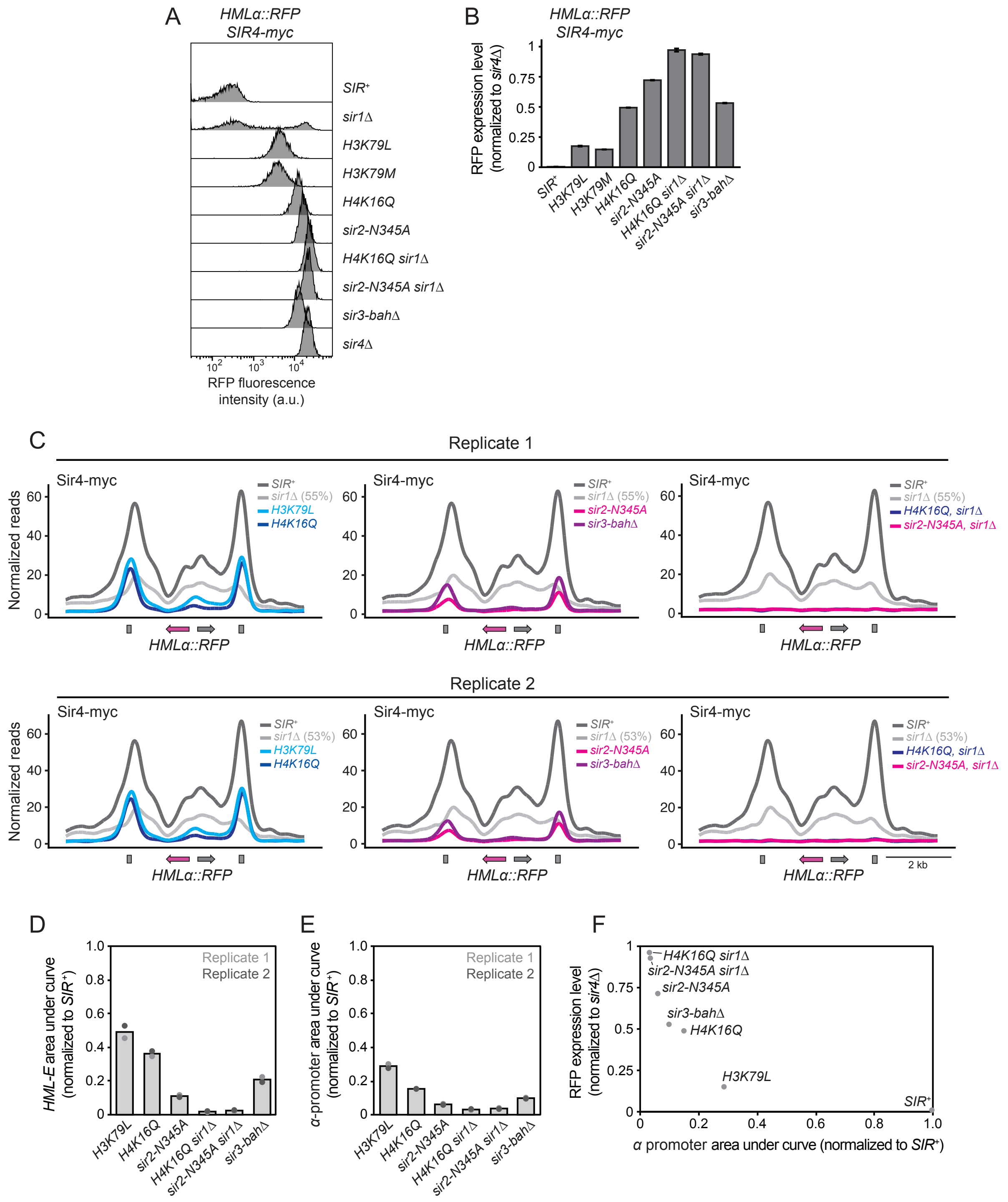


**Figure S6: Spreading mutants affected Sir4 nucleation at silencers in *HMLα::RFP*.** A. Fluorescence intensity profiles for different spreading mutants and control strains (JRY11494, JRY13129, JRY13131, JRY13241-13244, JRY13248, JRY13249, JRY13633). All strains contained *HMLα::RFP* and *SIR4-myc*. Strains were grown and analyzed in biological triplicate, and representative flow cytometry profiles are shown. B. Quantification of RFP expression levels from A, as calculated via the geometric mean intensity statistic in FlowJo. Data are means ± SD (n = 3 biological replicates). C. ChIP of Sir4-myc in different spreading mutant backgrounds, with two biological replicates. Reads were normalized to the genome-wide median. The percent of silenced cells in *sir1Δ* is shown next to the *sir1Δ* label. D-E. Quantification of Sir4-myc ChIP in spreading mutants. The area under curve was calculated at *HML-E* (D) and the $\alpha$ promoter (E) for data shown in C. Gray bars represent the mean of both replicates for each strain. H. Relationship between the RFP expression level (B) and amount of Sir4-myc at the $\alpha$ promoter (E).

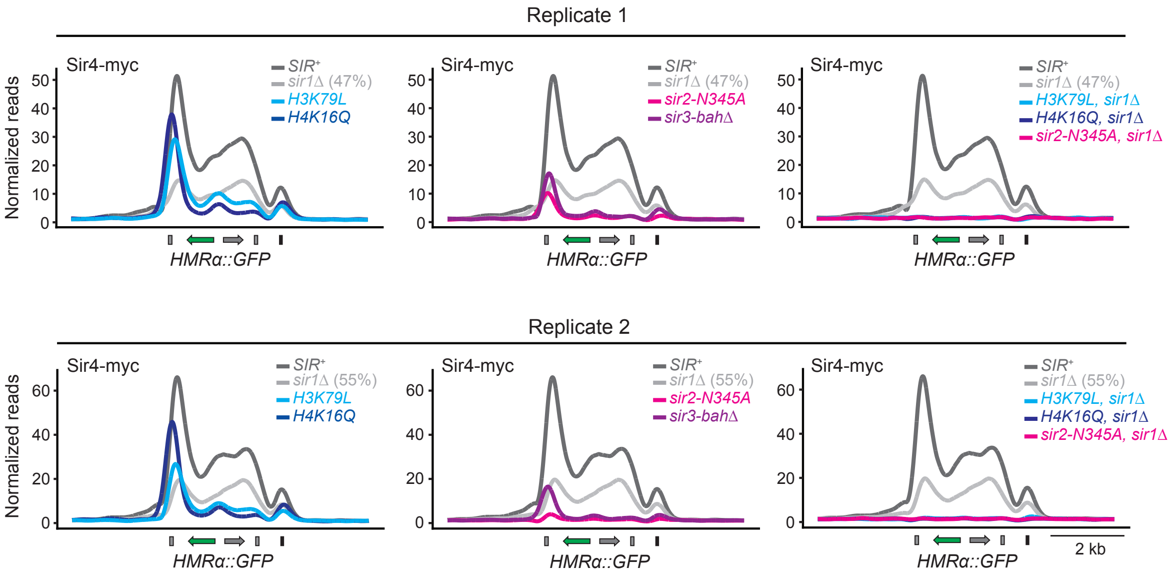


**Figure S7: Biological replicates of Sir4-myc ChIP in spreading mutants.** Reads were normalized to the genome-wide median. Replicate 1 is identical to Figure 5C-E and is shown here for convenience (JRY13130, JRY13132, JRY13233, JRY13234, JRY13236, JRY13237, JRY13239, JRY13631, and JRY13632). The percent of silenced cells in *sir1Δ* is shown next to the *sir1Δ* label. Quantification for both replicates is shown in Figure 5F-G.

| Condition | Silenced (S) | Expressed (E) | E → S ($k_S$, gen$^{-1}$) | S → E ($k_E$, gen$^{-1}$) | S/(S+E) | $k_S/(k_S+k_E)$ |
|---|---|---|---|---|---|---|
| *sir1Δ* 100 nM | 12 | 88 | 0.02 | 0.16 | 0.12 | 0.12 |
| *sir1Δ* 125 nM | 25 | 75 | 0.04 | 0.10 | 0.25 | 0.27 |
| *sir1Δ* 175 nM | 56 | 44 | 0.09 | 0.03 | 0.56 | 0.74 |
| *sir1Δ* 250 nM | 89 | 11 | 0.19 | 0.02 | 0.89 | 0.91 |
| *SIR1* 40 nM | 18 | 82 | 0.02 | 0.30 | 0.18 | 0.07 |
| *SIR1* 50 nM | 46 | 54 | 0.05 | 0.16 | 0.46 | 0.24 |
| *SIR1* 70 nM | 61 | 39 | 0.13 | 0.10 | 0.61 | 0.57 |
| *SIR1* 100 nM | 83 | 17 | 0.23 | 0.03 | 0.83 | 0.87 |

**Table S1: Frequencies of different epigenetic states and switching rates at different Sir4 dosages.** Data were collected from live-cell microscopy of different conditions over a 10-hr time course (JRY13092 and JRY13093). The percentages of cells exhibiting the silenced state (S) and expressed state (E) were calculated by manually counting cells in each state at 5 hrs (n > 500 cells per condition). The switching rates were calculated from the same time course and are shown in Figure 2G-H. $k_S$ reflects the rate of silencing establishment, and $k_E$ reflects the rate of silencing loss. If the $k_S$ and $k_E$ rates can account for the observed frequencies of silenced and expressed cells, then S/(S+E) should be similar to $k_S/(k_S+k_E)$, as observed.

**Table S2: Strains and oligonucleotides used in this study**

**Movie S1: Live-cell microscopy of *sir1Δ HMRα::GFP LexA-ER-AD LexO-SIR4* cells at 125 nM estradiol.** Both GFP and brightfield images are shown (JRY13093).

**Movie S2: Live-cell microscopy of *SIR1 HMRα::GFP LexA-ER-AD LexO-SIR4* cells at 50 nM estradiol.** Both GFP and brightfield images are shown (JRY13092).

**Movie S3: Time-lapse movie of *HMRα::RFP/HMRα::GFP KAR1/kar1-1 SIR1* heterokaryons.** *HMRα::RFP KAR1 SIR1* (JRY13727) haploids were mixed with *HMRα::GFP kar1-1 SIR1* haploids (JRY13708) in 20 nM estradiol. A small subset of these cells mated to form heterokaryons, which have a characteristic bi-lobed shape. These heterokaryons produce haploid progeny that each contain a nucleus derived from one of the two heterokaryon nuclei (Conde and Fink, 1976). Each example shown has a heterokaryon that either maintains a silenced state at one allele but not at the other or experiences a switching event at one allele but not the other. Each row is a field of view with one or more such heterokaryons. The first column shows brightfield, GFP, and RFP channels. The second column shows GFP and RFP channels. The third column shows only the GFP channel, and the fourth column shows only the RFP channel. The timestamp is shown in the upper left, and a scale bar is shown in the upper right (5 µm).
